# Advances in genomic characterization of *Urochloa humidicola*: exploring polyploid inheritance and apomixis

**DOI:** 10.1101/2023.08.31.555743

**Authors:** Aline da Costa Lima Moraes, Marcelo Mollinari, Rebecca Caroline Ulbricht Ferreira, Alexandre Aono, Letícia Aparecida de Castro Lara, Marco Pessoa-Filho, Sanzio Carvalho Lima Barrios, Antonio Augusto Franco Garcia, Cacilda Borges do Valle, Anete Pereira de Souza, Bianca Baccili Zanotto Vigna

## Abstract

Tropical forage grasses are an important food source for animal feeding, with *Urochloa humidicol*a, also known as Koronivia grass, being one of the main pasture grasses for poorly drained soils in the tropics. However, genetic and genomic resources for this species are lacking due to its genomic complexity, including high heterozygosity, evidence of segmental allopolyploidy, and reproduction by apomixis. These complexities hinder the application of marker-assisted selection (MAS) in breeding programs. Here, we developed the highest-density linkage map currently available for the hexaploid tropical forage grass *U. humidicola*. This map was constructed using a biparental F1 population generated from a cross between the female parent H031 (CIAT 26146), the only known sexual genotype for the species, and the apomictic male parent H016 (BRS cv. Tupi). The linkage analysis included 4,873 single nucleotide polymorphism (SNP) markers with allele dosage information. It allowed mapping of the apospory locus and phenotype to linkage group 3, in a region syntenic with chromosome 3 of *Urochloa ruziziensis* and chromosome 1 of *Setaria italica*. We also identified hexaploid haplotypes for all individuals, assessed the meiotic configuration, and estimated the level of preferential pairing in parents during the meiotic process, which revealed the autopolyploid origin of sexual H031 in contrast to H016, which presented allopolyploid behavior in preferential pairing analysis. These results provide new information regarding the genetic organization, mode of reproduction, and allopolyploid origin of *U. humidicola*, potential SNPs markers associated to apomixes for MAS and resources for research on polyploids and tropical forage grasses.

**Key message:** We present the highest-density genetic map for the hexaploid *Urochloa humidicola*. SNP markers expose genetic organization, reproduction, and species origin, aiding polyploid and tropical forage research.

## 1 Introduction

Tropical forage grasses greatly impact the world’s economy, as they are the main food source for animals in tropical and subtropical regions (Pereira et al. 2018a; Simeão et al. 2021). The African species *Urochloa humidicola* (Rendle) Morrone & Zuloaga (syn. *Brachiaria humidicola* (Rendle) Schweick.), commonly known as Koronivia grass, is one of the main pasture grasses cultivated in these regions (Ferreira et al. 2021). Despite its economic importance, *U. humidicola* has few available genetic and genomic resources, mainly due to its genomic complexity, including a high ploidy level (2n=6x=36, 48 or 54), evidence of segmental allopolyploidy and high heterozygosity. In addition, the species reproduces predominantly by apomixis (Boldrini et al. 2009; Vigna et al. 2016), an asexual reproductive mode in which the ovules forego meiosis and form seeds without fertilization, resulting in genetic replicas (clones) of the female parent. Consequently, the application of marker-assisted selection (MAS) is still limited in *U. humidicola* breeding programs.

Advances in large-scale genotyping technologies and genotypic software have allowed the identification of many high-quality single nucleotide polymorphisms (SNPs) with allele dosage information in tropical forage grasses (Bourke et al. 2018a; Grandke et al. 2017; Mollinari and Garcia 2019). In this context, robust genomic studies have been recently reported, including genome-wide association studies (Matias et al. 2019a), genomic predictions (Aono et al. 2022; de C Lara et al. 2019; Martins et al. 2021; Matias et al. 2019b), and genetic maps (Deo et al. 2020; Ferreira et al. 2019; Worthington et al. 2016, 2019, 2021). Additionally, the recent assembly of two diploid genomes of *Urochloa ruziziensis* (Pessoa-Filho et al. 2019; Worthington et al. 2021) provided an invaluable resource for progress in genomic studies and molecular breeding of *Urochloa* grasses (Ferreira et al. 2021).

The construction of genetic maps, used for understanding the genetic organization of a species, is challenging in polyploids, mainly due to the wide range of possible meiotic configurations (Mollinari et al. 2020). In addition, the genomic complexity and self-incompatibility of such plants explain few genetic maps for *Urochloa* species available. The majority of genetic maps in polyploids were constructed using two-point or pairwise methodologies. This approach uses isolated marker pairs to detect recombination events and was originally developed to encompass simplex or single-dose markers, which segregate in a 1:1 proportion (Wu et al., 1992). The advent of high-throughput genotyping technologies has enabled the assessment of markers exhibiting variation across multiple homologs, referred to as multiple-dose or multiplex markers (Bourke et al. 2018b; Grandke et al. 2017; Hackett et al. 2017).

Despite the computational efficiency of two-point methods, they cannot exploit the information of multiple markers simultaneously. This limitation is especially significant in polyploid species, where datasets typically present a low signal-to-noise ratio. To overcome this limitation, multilocus methods have been developed for tetraploids (Leach et al. 2010; Zheng et al. 2021) and plants with higher ploidy levels (Mollinari and Garcia 2019). The latter approach is implemented in an R package called MAPpoly. These methodological advancements have increased the potential use of multiple-dose markers in assessing intricate polyploid inheritance systems, thereby facilitating the development of considerably more robust and informative genetic maps (Mollinari et al. 2020; Oloka et al. 2021).

To date, two genetic maps have been published for *U. humidicola* (Vigna et al. 2016; Worthington et al. 2019), both constructed using pipelines developed for diploid organisms and without accounting for possible allele dosages, polysomic segregation, and multilocus information, which would enable more robust inference in a hexaploid species. Therefore, generating a multilocus genetic map is crucial for systematically characterizing the inheritance patterns in this species. In addition, preferential pairing profiles, which may be inferred from genetic maps, can help test hypotheses about a species’ origin and are also relevant for the study of genome evolution (Kamiri et al. 2018; Okada et al. 2010).

Another useful application of genetic maps is the mapping of loci related to target traits, such as the mode of reproduction. The type of apomixis in the *Urochloa* genus is apospory (Ferreira et al. 2021), which is characterized by the differentiation of adjacent nucellar cells into unreduced embryo sacs, a process named apomeiosis, followed by the development of the unreduced egg into an embryo without fertilization (parthenogenesis) (Xiong et al. 2023). In Paniceae grasses, the components of parthenogenesis are usually inherited together as a single dominant Mendelian factor with suppressed recombination and denoted as the “apospory-specific genomic region” (ASGR) (Kaushal et al. 2019; Palumbo et al. 2022).

The sequence of the ASGR-BABY BOOM-like (ASGR-BBML) gene, proposed as a candidate for parthenogenesis in *Cenchrus ciliaris*/*Pennisetum squamulatum*, was used to develop the psASGR–BBML-specific primer pair p779/p780, which has already been validated in some grasses (Akiyama et al. 2011; Worthington et al. 2016, 2019). Recently, the specific amplicon from p779/p780 was observed to be in full linkage with the ASGR in an F_1_ mapping population of *U. humidicola* (Worthington et al. 2019). However, mapping this marker in other populations is important to verify candidate genes for the parthenogenesis component of apomixis. In addition, such mapping will allow a better understanding of the molecular basis and inheritance of apomixis, assisting in practical applications for modern agriculture.

To the best of our knowledge, we are presenting the highest-density genetic map available for the hexaploid *U. humidicola*, generated with SNP markers with allele dosage information and using a multilocus approach. This map was used to identify a region linked to apomixis, assess collinearity with the related species *U. ruziziensis*, evaluate preferential pairing profiles during meiosis in the parents, construct haplotypes, and estimate the meiotic configuration for all individuals of a biparental F_1_ population. In addition to providing new information regarding the genetic organization, mode of reproduction and allopolyploid origin of *U. humidicola*, these results represent new advances and resources for tropical forage grasses and research communities that study polyploids.

## 2 Material and methods

### 2.1 DNA extraction, GBS library preparation and sequencing

A full-sib progeny set consisting of 279 F_1_ hybrids was obtained from a cross between the hexaploid apomictic pollen donor *U. humidicola* cv. BRS Tupi (hereafter H016) and the hexaploid sexual accession BRA005811-H031 (CIAT 26146, hereafter H031), as described by Vigna et al. (2016). These intraspecific progeny are part of the *Urochloa* breeding program of Embrapa Gado de Corte (Brazilian Agricultural Research Corporation), located in Campo Grande/MS, Brazil (20°27′ S, 54°37 56′ W, 530 m). Leaf samples were collected from each hybrid and its parents and subjected to genomic DNA extraction according to Doyle and Doyle (1987). DNA concentration and quality were examined by electrophoresis on a 2% (w/v) agarose gel and a Qubit 3.0 fluorometer (Thermo Scientific, Wilmington, USA), respectively.

A DNA genotyping-by-sequencing (GBS) library was constructed for all F_1_ individuals (one sample each) and the two parental genotypes (five replicates each) of the mapping population. Genomic DNA (210 ng per individual) was processed using a combination of a rarely cutting enzyme (*Pst*I) and a frequently cutting enzyme (*Msp*I), as described by Poland et al. (2012). Libraries were sequenced as 150-bp single-end reads on the NextSeq 500 platform (Illumina, San Diego, CA, USA), and the quality of the resulting sequence data was evaluated using the FastQC toolkit (Patel and Jain, 2012).

### 2.2 GBS SNP calling and allele dosage estimation

SNP calling was performed using the Tassel-GBS pipeline (Glaubitz et al., 2014) modified to obtain the read counts for each SNP allele (Pereira et al. 2018b). A previous study (Martins et al. 2021) revealed a few putative apomictic clones of the female parent in a biparental population. Therefore, 62 hybrids were removed, and the next analyses were performed using 217 hybrids. GBS tags were aligned against the *U. ruziziensis* genome (Pessoa-Filho et al. 2019; https://www.ncbi.nlm.nih.gov/data-hub/genome/GCA_015476505.1/), the most closely related reference genome available. The Bowtie2 algorithm version 2.1 (Langmead and Salzberg, 2012) was used to align reads using the following settings: a limit of 20 dynamic programming problems (D), a maximum of 4 times to align a read (R), and the very-sensitive-local argument.

SuperMASSA software (Serang et al. 2012) was used to estimate the probabilities of allele dosages for each individual and SNP combination. Data points with a maximum probability lower than 0.75 were assumed missing. The minimum average read depth considered was 20 reads, and the model used was the F_1_ population model. Markers were fitted to ploidies 2, 4, and 6, and those classified as ploidy six were selected. A second round of quality filtering was conducted to filter out SNPs with fewer than 30 reads on average across all SNPs and markers with redundant information. The resulting dataset was imported into the software MAPpoly (Mollinari and Garcia 2019; Mollinari et al. 2020), available at https://CRAN.R-project.org/package=mappoly). SNPs with more than 25% missing data were removed using the function “filter_missing.” Finally, a chi-square test of goodness-of-fit to expected Mendelian segregation ratios was performed. SNPs displaying P < 2.5 x 10^-6^ (computed through Bonferroni’s correction) were filtered out using the function “filter_segregation”.

### 2.3 Linkage map construction

The genetic map was constructed using the R package MAPpoly (Mollinari and Garcia 2019; Mollinari et al. 2020). First, we estimated the pairwise recombination fraction between all screened SNPs and selected the most likely linkage phase configurations for each pair. Next, we assembled a recombination fraction matrix using the *U. ruziziensis* genome order. Although we observed linkage blocks, it was not possible to cluster them into linkage groups (LGs) due to the low signal-to-noise ratio typically observed in hexaploid species. To cope with the missing information, we aggregated the recombination fraction and logarithm of odd (LOD) scores of neighboring SNPs by averaging cells in a grid of 10 x 10 SNPs in a lower-resolution matrix, resulting in more apparent linkage blocks.

Then, we applied clustering analyses using the unweighted pair group method with arithmetic mean (UPGMA) algorithm to generate SNP clusters corresponding to the LGs. We applied the ‘rf_snp_filter’ function from MAPpoly to each LG, eliminating SNPs that failed to meet an LOD score threshold of 5.0 for both phase configuration and linkage. This was achieved with respective filtering quantiles of 0.05 and 0.8. Subsequently, we applied the multidimensional scaling (MDS) algorithm (Preedy & Hackett, 2016) to the low-resolution recombination fraction matrix. After a visual inspection, we removed blocks disrupting the monotonicity of the matrix. Given the MDS-based order, we removed the SNP block restriction, yielding the order of the SNPs used in the map.

We re-estimated the linkage phases and recombination fractions for each variant in all LGs using the algorithms implemented in MAPpoly’s function “est_rf_hmm_sequential”. The algorithm commences by exhaustively searching for the best phase configuration for a small subset of five markers at the beginning of the LG using the multilocus likelihood based on the hidden Markov model (HMM) approach (Mollinari and Garcia 2019). Thereafter, the remaining markers are inserted sequentially using the pairwise linkage information to phase the allelic variants in parental homologs based on their LOD score. In the event of multiple-phase alternatives, the multilocus likelihood of the map is used to select the best phase configuration for the next round of marker insertion. Markers that inflate the map over a threshold of 3 centimorgans (cM) or yield more than 20 linkage phase configurations are not inserted into the map. This process is repeated until all markers are positioned in the phased map. Finally, the genetic map is reconstructed while considering a global genotyping error of 10%.

### 2.4 Comparative analysis

Synteny between the *U. humidicola* genetic map and *Setaria italica* (foxtail millet) genome was carried out by anchoring mapped SNPs into the *U. ruziziensis* genome. In brief, sequences flanking SNPs in the present map of *U. humidicola* were extended to a length of 2000 bp (1000 bp to each side) using the position in the reference genome of *U. ruziziensis* (Pessoa-Filho et al. 2019). Then, these extended sequences were queried against the chromosomes of the *S.italica* genome retrieved from the Phytozome database v.13 using Basic Local Alignment Search Tool (BLAST) with a cutoff E-value of < 1 × 10^−5^. The resulting synteny between each *U. humidicola* LG and the chromosomes of *S. italica* was illustrated using the ggplot2 R package (Goodstein et al. 2012). LGs of *U. humidicola* were plotted using cM lengths, while chromosomes of *S. italica* were plotted using physical lengths.

### 2.5 Preferential pairing profile

We used the methodology described by Mollinari et al. (2020) to assess the preferential pairing that may occur between all homologs of each parent during hexaploid meiosis in *U. humidicola*. Briefly, we estimated the posterior probability distribution of each of the 15 possible pairing configurations at any position in the genome for both parents, assessing the preferential pairing for specific homolog pairs. To test whether the observed homolog configurations differed from their expected frequencies under random pairing, we used the χ^2^ test with < 10^−4^ to declare significance.

### 2.6 Haplotype reconstruction and meiotic configuration assessment

Offspring haplotype reconstruction and the meiotic configuration assessment were performed using the methodology proposed by Mollinari et al (2020). Briefly, we estimated the probability that an offspring carried a particular genotype at a specific position. Then, we combined this information to build six profiles indicating the probability of inheritance of a particular homolog across whole chromosomes for all offspring and both parents in the *U. humidicola* mapping population. Using this information, we detected the position and the homologs involved in recombination events across the genome of all offspring. Then, we classified the recombination events according to the number of homologs involved in the recombination chain: if two or fewer homologs were involved, we considered it evidence of bivalent pairing; if three or more homologs were involved, we considered it evidence of multivalent pairing.

### 2.7 Genotyping with the ASGR-BBML-specific primer pair p779/p780 and amplicon sequencing

The parents and progeny of the mapping population were analyzed with p779/p780, a primer pair specific to the candidate gene for the parthenogenesis component of apomixis that has been developed for *Pennisetum squamulatum* (primer sequence: TATGTCACGACAAGAATATG, TGTAACCATAACTCTCAGCT; Akiyama et al. 2011) and validated in *U. humidicola* (Worthington et al. 2019). Primers were used to amplify DNA under the following PCR conditions: each forward and reverse primer at 0.5 µM, 1X GoTaq® Colorless Master Mix (Promega), and 40 ng of DNA. Thermocycling was performed with an initial denaturation at 94°C for 5 min followed by 35 cycles of 94°C for 30 secs, 59°C for 30 secs and 72°C for 60 secs, with a final extension step at 72°C for 10 min. Amplified products (12 µl) for both markers were resolved on 1.5% agarose gels stained with ethidium bromide.

The p779/p780 primer pair was evaluated using Sanger technology to sequence the p779/780 amplicon of three random apomictic hybrids (named H261, H310 and H330), in addition to four replicates of the apomictic parental genotype H016. The PCR products were generated as previously described and purified by precipitation with polyethylene glycol (Schmitz and Riesner, 2006). The sequencing cycle was performed using the BigDye Terminator kit v. 3.1 and cleaned up with EtOH/EDTA (adapted from Moreau 2014). The purified products were sequenced on an ABI 3500xl Genetic Analyzer (Applied Biosystems). The resulting data were analyzed and filtered by quality parameters using Chromas software (http://www.technelysium.com.au/Chromas.html), and consensus sequences were generated using CAP3 (https://doua.prabi.fr/software/cap3). Singletons were annotated through blastn sequence comparison to the Nucleotide Collection (nr/nt) and then individually compared with *C. ciliaris* and *P. squamulatum* ASGR-BBM-like1 gene sequences using the Clustal Omega Multiple Sequence Alignment algorithm.

### 2.8 Apospory and p779/p780 marker mapping

In previous studies (do Valle et al. 2008; Zorzatto et al. 2010; de Araujo Bitencourt et al. 2012), the reproductive mode of each F1 individual was assessed by examining embryo sacs using the methodology described by Young et al. (1979). The phenotype (embryo sac evaluation) and the genetic marker p779/p780 were expected to indicate the mode of reproduction (apomictic vs. nonapomictic). Therefore, using binary classification, both markers were mapped onto the preconstructed map. First, pairwise linkage analysis was conducted between markers positioned on the map and the reproduction mode markers using MAPpoly. After identifying the chromosome where the markers were linked, their best position was determined by inserting them between all neighboring markers and selecting the map with the maximum multipoint likelihood.

## 3 Results

### 3.1 SNP calling and genetic map

After GBS library sequencing, FastQC revealed 1,727,547,916 high-quality sequencing reads, 1,065,791,025 of which were assigned to 73,685,381 tag pair sites using the Tassel-GBS pipeline for polyploids (Pereira et al. 2018b). Then, 105,539 SNPs were identified through alignment taxa of 33.91% with the *U. ruziziensis* reference genome. SuperMASSA software successfully estimated the ploidy and allele dosage of 20,390 markers.

We excluded the offspring Bh152, Bh181, Bh226, and Bh245 from the study owing to inadequate read depth, reducing the total number of individuals to 213. After quality filtering, 7,069 high-quality SNPs with allele dosage information were retained for later analyses (Supplementary Fig. 1). Of these SNPs, 54.2% were classified as simplex, 2.4% as double-simplex, and 43.5% as higher-dosage markers (multiplex).

The UPGMA algorithm resulted in six clusters representing the *U. humidicola* LGs (Supplementary Fig. 2), three corresponding to whole *U. ruziziensis* chromosomes and the others corresponding to combinations of multiple *U. ruziziensis* chromosomes. In the ordered recombination fraction matrix (Supplementary Fig. 3), we noticed a block-diagonal pattern showing the six LGs and monotonicity in all submatrices, indicating that the MDS algorithm provided a good global order for the marker blocks. The reference genome-assisted reordering yielded a genetic map with 3,821 unique SNPs and a set of redundant markers, resulting in a total of 4,873 SNP markers covering 654.44 cM, with LGs ranging from 75.12 cM to 131.65 cM in length and an average marker density of 7.45 markers/cM (Fig. 1, Table 1, and Supplementary Fig. 4). The LG with the most markers was LG6, which included 1,099 markers with a density of 9.73 markers/cM. Conversely, LG5 was the sparsest, having only 191 markers and the lowest marker density of 1.62 markers/cM. LG2 was the longest, extending a genetic length of 131.65 cM, and contained 955 markers with a density of 7.25 markers/cM. LG3 showed the highest marker density at 11.70 markers/cM, followed by LG6 with 9.73 markers/cM (Table 1). The intermarker distance was consistently less than one centimorgan across all LGs. Noteworthy gaps were present in LG5 (16.21 cM), LG1 (8.12 cM), and LG4 (7.76 cM). Although LG5 contained the fewest markers, it covered a length of 117.85 cM. Although this feature dilutes the group’s marker density, it ensures comprehensive coverage of the corresponding chromosome.

**Fig. 1.**
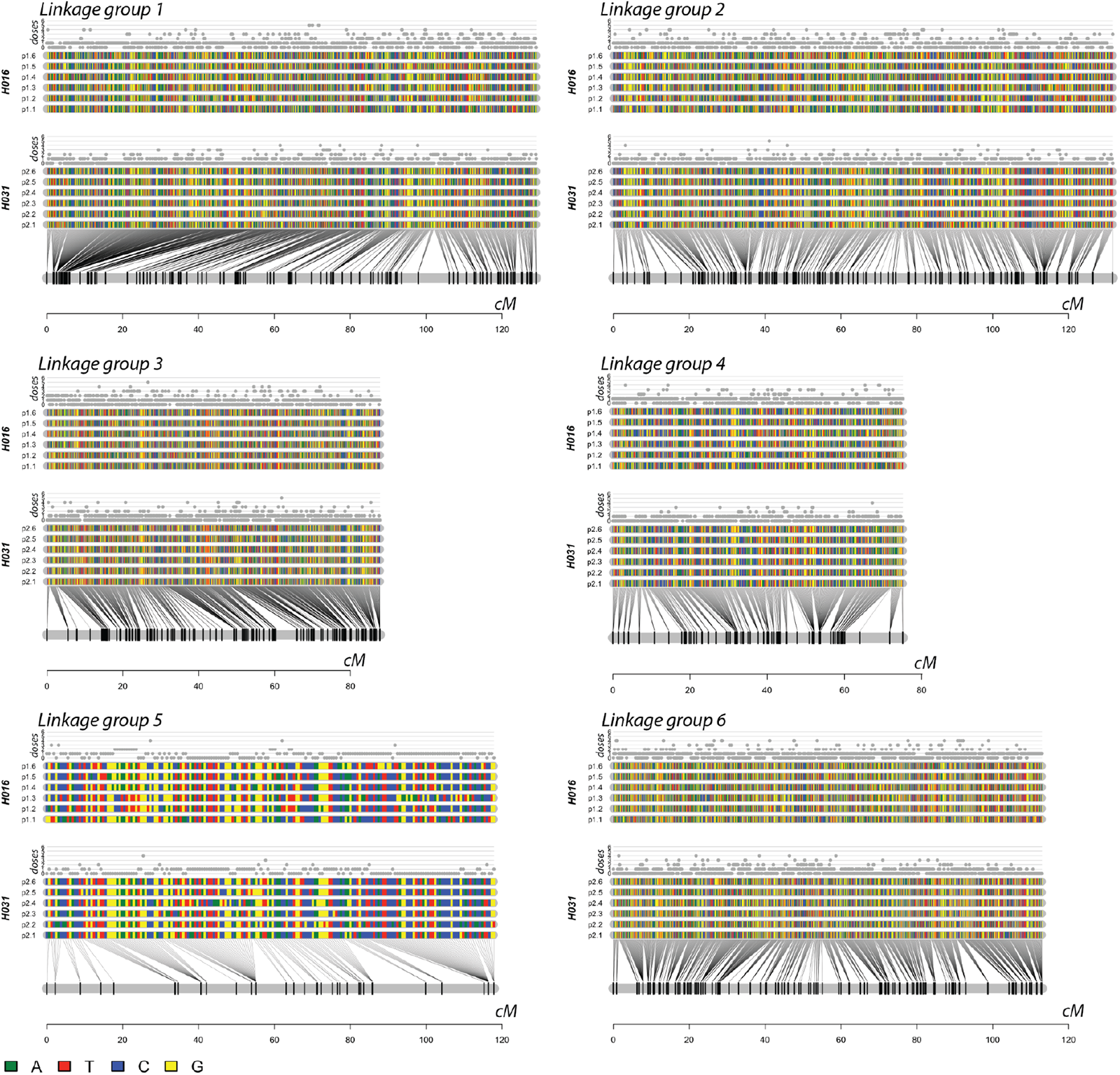
Genetic phased map of *U. humidicola* showing the parental linkage phase configuration of homology groups for the parental strains H016 and H031 for all six linkage groups. The colored rectangles indicate the configuration of the SNP dosages in the six homologs for H016 (p1.1 to p1.6) and in the six homologs for H031 (p2.1 to p2.6)

**Table 1.** Summary of the *U. humidicola* genetic linkage map.

| LG | Reference chromosome<sup>1</sup> | Map size (cM) | Markers/c M | Number of markers |  |  | Total | Maximum gap |
| --- | --- | --- | --- | --- | --- | --- | --- | --- |
|  |  |  |  | S<sup>2</sup> | DS<sup>3</sup> | MP<sup>4</sup> |  |  |
| 1 | 1 | 129.06 | 8.12 | 749 | 12 | 287 | 1048 | 8.12 |
| 2 | 2-9 | 131.65 | 7.25 | 689 | 20 | 246 | 955 | 5.12 |
| 3 | 3-6 | 87.79 | 11.7 | 744 | 13 | 270 | 1027 | 5.57 |
| 4 | 4 | 75.12 | 7.36 | 463 | 6 | 84 | 553 | 7.76 |
| 5 | 5 | 117.85 | 1.62 | 142 | 19 | 30 | 191 | 16.21 |
| 6 | 7-8 | 112.97 | 9.73 | 847 | 26 | 226 | 1099 | 5.84 |
| Total | - | 654.44 | 7.45 | 3634 | 96 | 1143 | 4873 | - |

<sup>1</sup>*U. ruziziensis* reference chromosome; <sup>2</sup>S: simplex; <sup>3</sup>DS: double-simplex; <sup>4</sup>MP: multiplex

### 3.2 Comparative analysis

In the comparison of physical distance in *U. ruziziensis* versus the final ordered genetic distance in the six LGs of *U. humidicola*, we observed that LGs 1 and 5 were highly collinear with *U. ruziziensis* chromosomes 1 and 5, respectively (Supplementary Fig. 5). Nonetheless, three pairs of *U. ruziziensis* chromosomes (chromosomes 2 and 9; 3 and 6; and 7 and 8) were fused in three *U. humidicola* LGs (LG2, 3 and 6, respectively), and LG4 of *U. humidicola* presented an inversion when compared to chromosome 4 of *U. ruziziensis*, as well as the chromosome arms of LG 6 of *U. humidicola* when compared to chromosome 7 of *U. ruziziensis*. This comparative analysis demonstrated that the base chromosome number of *U. humidicola* is x = 6.

Then, synteny between the *U. humidicola* genetic map and *Setaria italica* (foxtail millet) genome was assessed by anchoring the 3,821 mapped SNPs into the *U. ruziziensis* genome. From the anchoring, each position in the reference genome was extended 1,000 bp to each side, resulting in 3,821 tags of 2,001 bp. These tags, used in a blastn analysis, resulted in 1,387 tags aligned against the *S. italica* genome. High collinearity was observed between LG5 of the *U. humidicola* genetic map and chromosome 8 of the *S. italica* genome (Supplementary Fig. 6).

Although some deviations were observed, LG1 and LG4 were also collinear with chromosomes 9 and 3. On the other hand, three pairs of *S. italica* chromosomes were fused in the *U. humidicola* genome: LG2 consisted of *S. italica* chromosomes 2 (chromosome arms) and 4 (centromere); LG3 consisted of chromosomes 1 (chromosome arms) and 7 (centromere); and LG6 consisted of chromosomes 5 (chromosome arms) and 6 (centromere). We observed that the chromosome arm regions of LGs 3 and 6 were inverted compared to those of the collinear chromosomes of *S. italica*.

### 3.3 Preferential pairing

Considering that in a hexaploid individual, there are 15 possible pairing configurations during prophase I of meiosis, we calculated preferential pairing on the basis of the probability profile for each of these possibilities. As shown in Fig. 2, the apomictic H016 genotype showed significant preferential pairing between homologs and at least one pair of homologs for all LGs. In contrast, we did not observe any significant level of preferential pairing across any homologs of the female parent, H031.

**Fig. 2.**
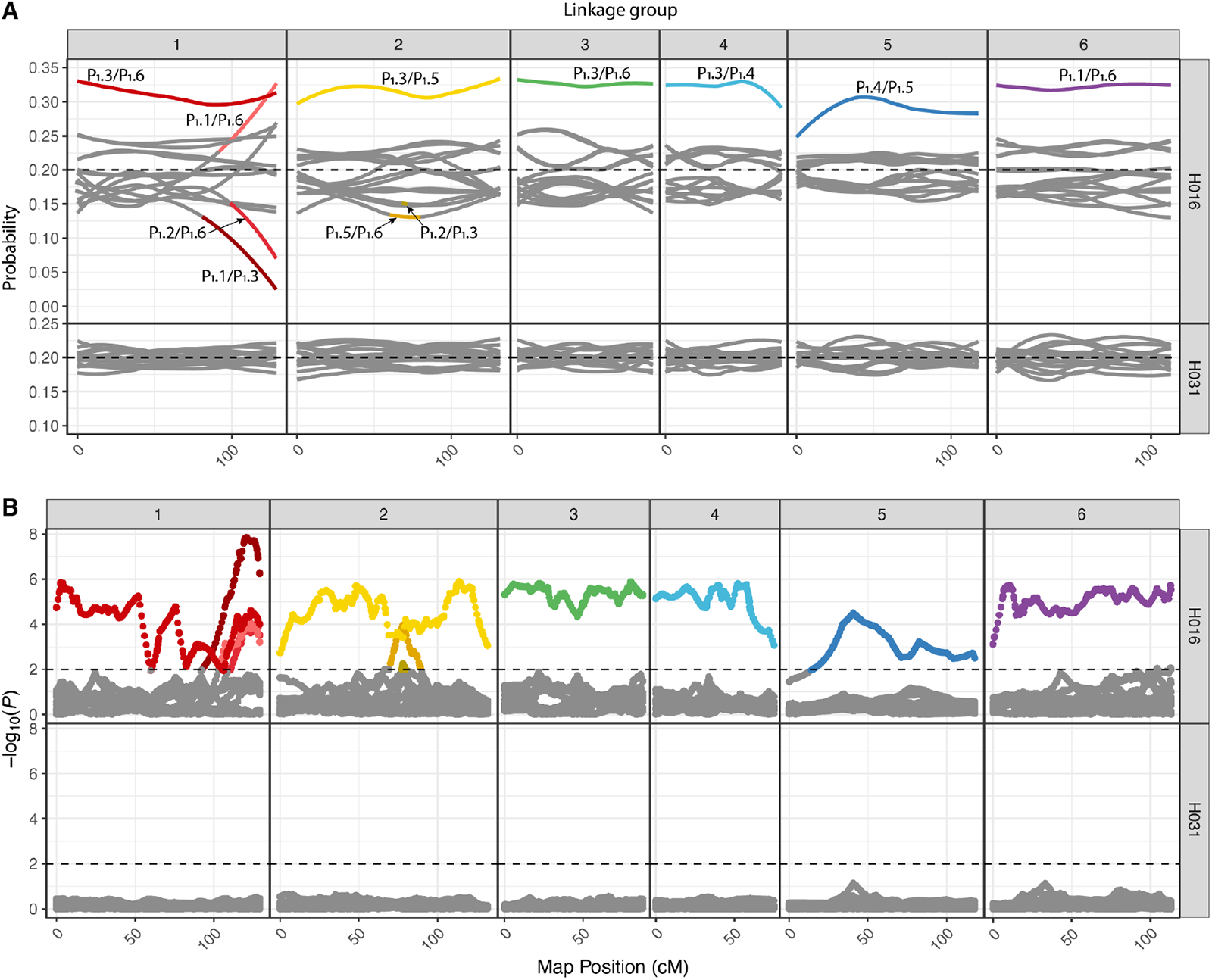
Preferential pairing profile in *U. humidicola*. (a) Probability profiles for 15 homolog pairs in the parents H016 and H031 across 6 LGs. The dashed lines specify the pairing probability expected under random pairing (3/15=0.2). (b) −log_10_(*P*) of a χ^2^ independence test for all possible homolog pairs, where dashed lines indicate *P* <10^−2^. The notation P_1_.X/P_1_.Y indicates the pairing configuration between homologous X and Y in the first parent, H016, for this particular instance. No significant level of preferential pairing was observed across the H031 sexual parent genome, but all LGs corresponding to the parent H016 showed a significant level of preferential pairing for at least one pair of homologs

### 3.4 Haplotype reconstruction and meiotic configuration assessment

From the parental phased chromosome information, we reconstructed the haplotype composition for each individual in the mapping population (Supplementary Material 1). Using the conditional probability distribution of the genotypes, we created 6 profiles (one for each homolog) representing the probability of inheritance of a specific homolog across the entire chromosomes for all individuals. With a heuristic procedure, we detected the crossing-over points between homologs in all hybrid individuals and assembled recombination chains that gave rise to their gametes, assessing the homologs involved in each meiosis event (Supplementary Fig. 7 shows an example for individual Bh1). Although some haplotype profiles exhibited inconsistent behavior across whole chromosomes, the percentage of inconclusive meiotic assessments for parents H016 and H031 was relatively low, at 5.5% and 7.7%, respectively.

We observed evidence of recombination chains involving a maximum of two homologs in 81.8% of meiosis events in H016 and 87.4% in the sexual parent H031, and the remaining 18.2% and 12.6% of events presented more than two recombinant homologs (Fig. 3). The presence of homologs in a recombination chain during meiosis suggests the formation of bivalent or multivalent structures. Bivalents are formed when two homologs are involved, while the involvement of more than two homologs suggests the formation of multivalents. Therefore, *U. humidicola* meiosis presents overall bivalent pairing bias rather than multivalent pairing, although these numbers may vary across LGs (Mollinari et al. 2020; da Silva Pereira et al. 2021). For instance, LG1 presented a multiple-chromosome chain percentage of 31.2% in the apomictic H016 genotype, while in LG4 of the sexual parent H031, this number was 3.8%.

**Fig. 3.**
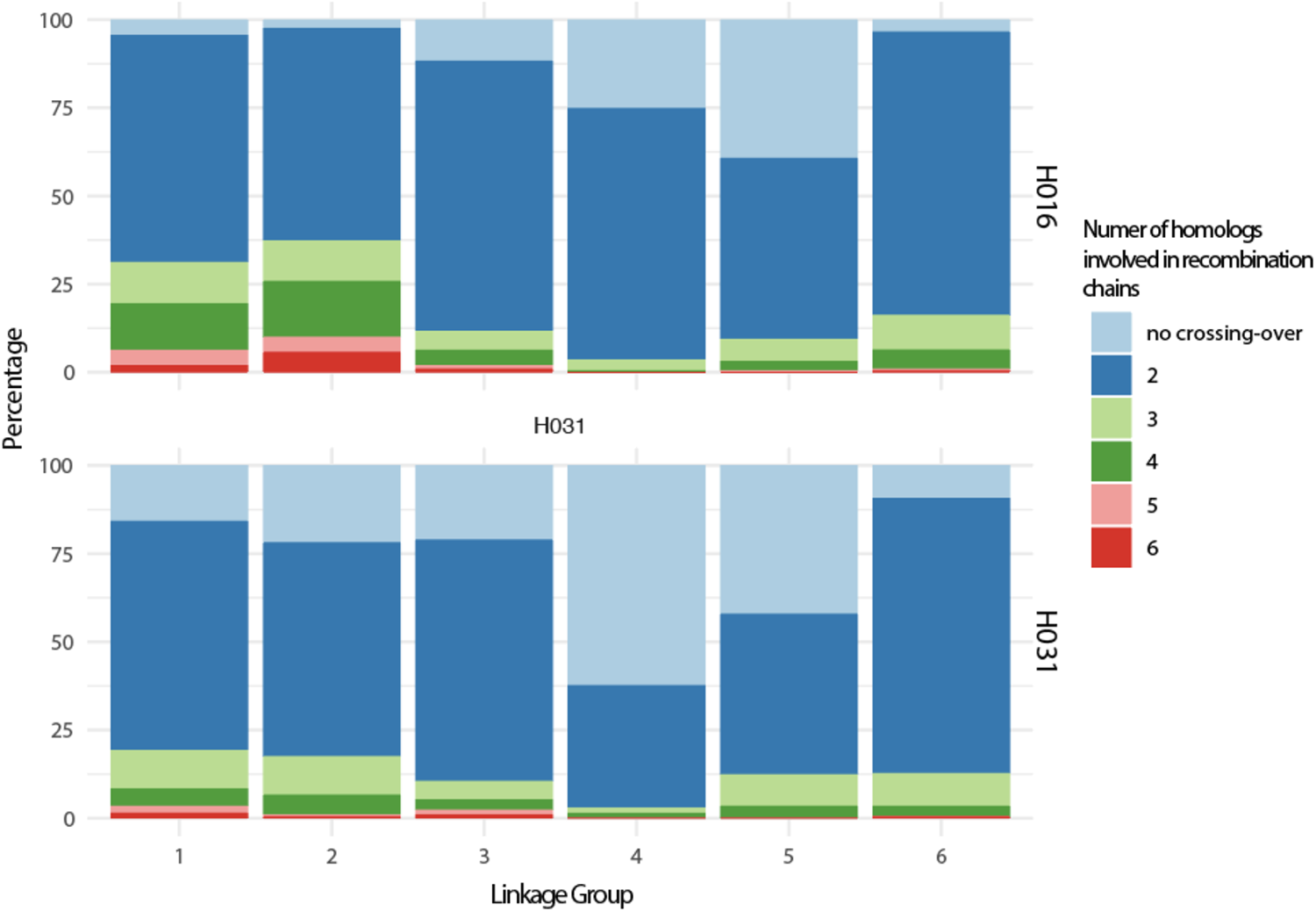
Percentage of cases where homologs do not recombine (no c.o., light blue) or recombine involving two (blue), three (light green), four (green), five (light rose) or six (red) homologs during metaphase I in the apomictic parent H016 and sexual parent H031 for all 6 LGs. Two homologs recombining suggests the presence of bivalents, while more than two suggests multivalent formation

### 3.5 Apospory mapping, genotyping with the ASGR-BBML-specific primer pair p779/p780 and amplicon sequencing

The ASGR-specific marker p779/p780 was assigned to position 53.73 cM of LG3, and the apomixis phenotype was assigned to position 47.16 cM of the same group (Fig. 4). A total of 12 SNP markers were in perfect linkage with the marker p779/p780, while five SNP markers were in linkage with the phenotype (Supplementary Fig. 8). Although LG3 is a composite of *U. ruziziensis* chromosomes 3 and 6, the p779/p780 marker and apomixis phenotype were mapped to a region that is clearly syntenic with *U. ruziziensis* chromosome 3. In addition, although the comparative analysis performed with *S. italica* showed that *U. humidicola* LG3 is a composite of *S. italica* chromosomes 1 and 7, the p779/p780 marker and the phenotype were mapped to a region syntenic with chromosome 1 of the *S. italica* genome.

**Fig. 4.**
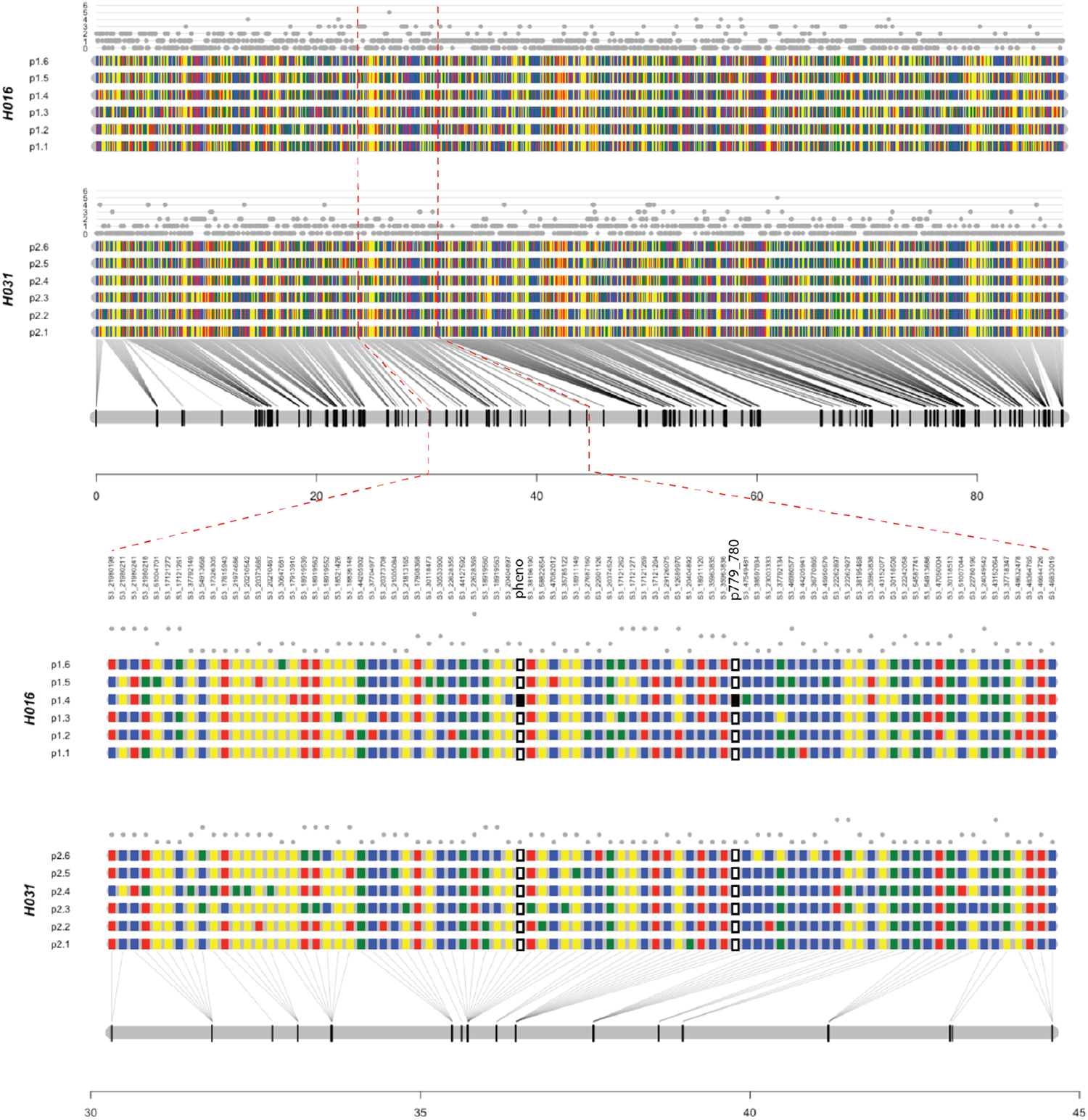
Linkage analysis of LG3 showing 10 SNP markers in linkage with the marker p779/p780, while 13 SNP markers were in linkage with the phenotype

The ASGR–BBML-specific marker p779/p780 was evaluated in the parents and hybrids of the *U. humidicola* population, and Supplementary Table S1 contains the genotype scores. Consensus singleton sequences were generated and compared to nr/nt, and the results showed significant blastn matches (E = 10-10) with ASGR-BBM-like1 gene sequences in *C. ciliaris* and *P. squamulatum*. Then, the singleton sequences were aligned with these gene sequences, and the multiple alignment revealed insertions and deletions (indels), in addition to SNPs (Supplementary Alignment 1).

## 4 Discussion

Climate change and the expected growth of the world’s population have raised concerns about food insecurity. Developing superior cultivars of tropical forage grasses can be effective and sustainable in supplying animal products, such as milk and beef (Pereira et al. 2018a; Simeão et al. 2021). To efficiently achieve these goals, we need to increase the amount of genetic and genomic information available for *Urochloa* spp., performing further studies on the genome composition and genetic architecture of target traits. This work presents the highest-density genetic map available for *U. humidicola*, a hexaploid tropical forage grass with evidence of segmental allopolyploid origin (Tomaszewska et al. 2023). We also constructed haplotypes for all offspring individuals, precisely mapped the apomixis locus, and estimated the level of preferential pairing in parents during meiosis, providing new insights into the origin of *U. humidicola*.

We used the GBS approach (Elshire et al. 2011; Poland et al. 2012) to genotype the *U. humidicola* F_1_ population, effectively capturing the genetic variation in this complex species with a large and highly heterozygous genome. Several statistical tools for marker dosage estimation have been developed, most assigning genotypic classes as discrete dosages (Blischak et al. 2018). Fortunately, some downstream applications can accommodate uncertainty in genotype calls (probabilistic genotypes), increasing the information available for each polyploid SNP locus (Gerard et al. 2018; Liao et al. 2021). The SuperMASSA tool (Serang et al. 2012) is one of these applications, and we used it to accurately assess the occurrence of all possible SNP dosages in *U. humidicola* for the first time.

Using all possible SNP dosage combinations rather than a subset to construct a genetic map allows for more accurate genotype inference, improves the estimation of recombination events, and provides greater resolution in identifying genetic variations (Soares et al. 2021).

These markers were used in linkage analysis for *U. humidicola*, resulting in a genetic map with 4,873 SNP markers, with 7.44 SNPs per cM, which to the best of our knowledge it is the most dense and informative genetic map published to date for the genus *Urochloa* (Ferreira et al. 2021, Worthington et al. 2019). For instance, the previous *U. humidicola* map was constructed using a smaller F1 population and molecular markers without allele dosage information, comprising separate linkage maps for each parent (Worthington et al. 2019). Despite the challenges posed by high ploidy, we successfully obtained a high-density genetic map for *U. humidicola* with six LGs and six homologs per LG, consistent with previous cytogenetic analyses (Jungmann et al. 2010; Penteado et al. 2000). In addition, our genetic map has a good pattern with defined visibility of the chromosome arms and centromere in almost all LGs (Fig. 1 and Supplementary Fig. 4), evidencing a high-quality linkage analysis.

Regarding the size of the genetic map, our map had a final haplotype size of 654.44 cM, and when multiplying it by the six chromosome sets, the total size would be 3926.64 cM, which is an intermediate size for both maternal (3558 cM) and paternal (4363 cM) maps obtained by Worthington et al. (2019). Furthermore, the genetic map size can be compared with *U. humidicola* genome sizes estimated by flow cytometry to obtain the genome-wide recombination rate (GWRR), a measure that provides information about the frequency of genetic recombination events along the entire genome and has been estimated for several species (Henderson et al, 2012). For H016, the observed genome size was 712.2 Mbp, resulting in a GWRR of 0.91 cM/Mbp, and for H031, with 725.35 Mbp, the rate was 0.90 cM/Mbp (Damasceno et al, 2023). This rate varies across different species, and even in genomic regions of the same species, and has important implications for genetic diversity, inheritance, and evolutionary processes.

We used the *U. ruziziensis* reference genome to improve the quality and robustness of our map, as well as to look for evidence of synteny, because this is the species with the closest phylogenetic relationship to *U. humidicola* for which whole-genome sequencing information is available (Pessoa-Filho et al. 2019). Assigning the *U. humidicola* LGs to *U. ruziziensis* chromosomes revealed high collinearity (Supplementary Fig. 5), which was expected because the two species were classified in different but sister taxa (Pessoa-Filho et al. 2017). Major disruptions were observed in LG2, LG3 and LG6, each of which split into two different chromosomes in *U. ruziziensis*, and in LG4, where we observed an inversion, evidencing some genomic rearrangements. These large structural differences compared to *U. ruziziensis* account for the lower base chromosome number of *U. humidicola* (x=6) and are consistent with the larger chromosome size found in three *U. humidicola* accessions than in other *Urochloa* species (Bernini and Marin-Morales 2001).

We also performed a comparative analysis with the *S. italica* genome, the second most closely related genome to that of *U. humidicola* available (Bennetzen et al. 2012). In contrast to those observed for *U. ruziziensis*, the marker positions in the *S. italica* genome do not exhibit strong linear correspondence with the *U. humidicola* map (Supplementary Fig. 6). However, a similar pattern of major disruptions was observed: LG2, LG3 and LG6 presented breakpoints, each corresponding to a different chromosome in *S. italica*, and LG4 presented an inversion compared to the corresponding *S. italica* chromosome. Using another version of the *U. ruziziensis* genome as a reference, Worthington et al. (2019) identified a similar pattern of collinearity between *S. italica* and *U. humidicola*: the same three pairs of *S. italica* chromosomes (1 and 7, 2 and 4, and 5 and 6) were fused compared to three LGs of *U. humidicola*. These findings verify the close evolutionary relationship between these species, suggesting that they share a common ancestor and have undergone similar patterns of chromosome evolution.

The inheritance of apospory in the F_1_ mapping population fitted the segregation ratio for a single dominant Mendelian factor, as previously observed in several Paniceae grasses (Ozias-Akins and van Dijk 2007), including *Megathyrsus maximus* (Deo et al, 2020) and *U. humidicola* (Vigna et al. 2016; Worthington et al. 2016, 2019; Zorzatto et al. 2010). The separate inheritance of apomeiosis and parthenogenesis has been confirmed for most apomictic model species, and apparently, these genetic apomixis elements are located in the same low-recombination region (Aguilera et al, 2015; Ortiz et al. 2020). In fact, both the phenotype and the ASGR locus were mapped to LG3 in this *U. humidicola* map, in a region syntenic with *U. ruziziensis* chromosome 3 and *S. italica* chromosome 1. Our results corroborate the observations made by Worthington et al. (2019), who also mapped the ASGR to a region syntenic with *S. italica* chromosome 1.

The ASGR locus was previously mapped to a region of reduced recombination in *Urochloa decumben*s LG5, syntenic with *S. italica* chromosome 5 (Worthington et al. 2016). On the other hand, in *U. ruziziensis* x *Urochloa brizantha* hybrids (Pessino et al, 1998), the apo locus was mapped to a region syntenic with rice chromosome 2 and maize chromosome 5, but both were syntenic to *S. italica* chromosome 1 (Zhang et al. 2012). For different species of *Paspalum*, the apo locus was mapped to a region syntenic with rice chromosome 12, and specifically for *Paspalum notatum*, to both chromosomes 12 and 2 (Ortiz et al. 2020). Therefore, although the genetic apomixis elements appear to be conserved among several species (Gualtieri et al. 2006), our results compared to those cited previously suggest that the apomixis-controlling loci occupy different positions in different genomes, as observed by other authors (Conner et al. 2008). In addition, this observation supports the hypothesis that apomixis emerged several times and independently during the evolution of the grass family (Ozias-Akins et al. 2003), making the identification of the genetic determinants of apomixis a challenge (Galla et al, 2019).

Genetic maps can also provide haplotype information, which is a combination of different polymorphisms with strong linkage disequilibrium (LD) carrying complete allele information about homolog inheritance across generations (Bhat et al. 2021). Although individual SNPs are useful for phenotype–genotype association studies, complex and quantitative traits, such as apomixis, may be regulated in polyploids by a set of SNPs or genes (haplotypes). Therefore, knowledge of allele configuration at a specific locus can be extremely useful for improving target traits in *U. humidicola*, as has already been shown in other important crops (Jensen et al. 2020; Mayer et al. 2020).

Several tools for polyploid haplotyping have been developed in recent years (Majidian et al. 2020; Zheng et al. 2021). From our map, we used the methodology proposed by Mollinari et al. (2020) to probabilistically recombine multiple SNPs and reconstruct haplotypes of all individuals in the mapping population. For the first time in *U. humidicola*, we assessed how the assembled parental homologs were transmitted to the hybrids, inferring the meiotic process. All LGs exhibited accurate information about parental haplotypes for most of their length. We observed most bivalent formation during meiosis in both parents, resulting in hybrids with two parental homologs, in addition to a small percentage of tri-, tetra-, penta-and hexavalents (Fig. 3). Our results corroborate the cytogenetics observation of bivalent and multivalent chromosome associations already reported in the meiosis of hybrids from the same *U. humidicola* mapping population (Vigna et al. 2016).

Although some authors reported the expectation of only multivalents in autopolyploid meiosis and only bivalents in allopolyploid meiosis (Bomblies 2023), the chromosome pairing behavior in polyploids can be more complex than originally thought (Lenormand et al. 2016; Zielinski and Mittelsten Scheid 2012). Several studies have shown that most established autopolyploid species form almost exclusively bivalents or, to a lesser degree, a mixture of bivalents and multivalents (Bomblies et al. 2016; Choudhary et al. 2020; Mason and Wendel 2020). On the other hand, multivalents were observed to some extent in allopolyploids (Zielinski and Mittelsten Scheid 2012). There is growing evidence that the occurrence of multivalents is indeed associated with newly formed polyploids (Mason and Wendel 2020). Previous work suggested that *U. humidicola* is a recent polyploid (Vigna et al. 2016), further corroborating the occurrence of multivalent formation during meiosis.

The haplotype analysis also revealed strong linkage disequilibrium (LD) between the apomixis phenotype and the p779/780 molecular marker on LG3, with a genetic distance of less than 7 cM. This confirms that the p779/780 marker exhibits low recombination with the apomixis trait, and can be used as a reliable genetic marker for detecting apomixis in *U. humidicola*. This marker has been successfully employed in Embrapa’s *Urochloa* spp. breeding program over the past few years, achieving a selection efficiency of 90%. We observed both the phenotype and the p779/780 marker carrying the same allelic variations (dosage 1 for the alternative allele) in the same homolog of the H016 genotype (homolog d in Fig. 4), while homozygosis for the reference allele was observed for the sexual parent at the same locus. The same allelic configuration was observed in two different SNPs that co-segregated with the phenotype. These SNPs exhibit significant potential for MAS upon validation, and may offer greater efficiency compared to the p779/780 marker. Our results corroborate the expected segregation observed in several species for the apomixis trait at a 1:1 ratio (presence:absence), which occurs in the progeny when the marker is present in one parent (for the apomixis trait, the apomictic H016 parental) yet absent in another parent (the sexual parent) (Wu et al, 1992).

The genetic map also allowed us to estimate the conditional probabilities of the offspring haplotypes and to uncover differences in homologous pairing profiles during meiosis, highlighting the power of saturated genetic maps to correctly diagnose preferential pairing (da Silva Pereira et al. 2021). While we did not observe any significant level of preferential pairing across the H031 genome, all LGs corresponding to the parent H016 showed preferential pairing for at least one pair of homologs (Fig. 2). Consequently, alleles present in different homologs of the apomictic parent H016 recombined with an equal chance, increasing the range of possible genotypes when compared to that in the sexual parent. In addition, our results suggest differences in genome composition between the sexual accession H031 and the apomictic accession H016.

These results corroborate previous linkage and in situ hybridization results that supported a segmental allopolyploid origin (AABBBB) and preferential pairing during meiosis in some apomictic accessions of *U. humidicola* but showed no evidence of subgenome differentiation in the sexual female parent H031 (BBBBBB) (Tomaszewska et al. 2023; Vigna et al. 2016; Worthington et al. 2019). In addition, two previous phylogenetic studies (Jungmann et al. 2010; Triviño et al. 2017) and a population structure study (Higgins et al. 2022) described a large genetic distance between H031 and most *U. humidicola* apomictic accessions of CIAT and EMBRAPA germplasm collections.

The H031 genotype is the only naturally occurring polyploid sexual genotype documented for *U. humidicola* and has considerable morphological differences when compared to other genotypes (Assis et al. 2014). Recently, this genotype was identified as an aneuploid with 2n=6x=36+1, a condition also previously reported for other accessions and interspecific hybrids of the genus *Urochloa* (Damasceno et al., 2023; Tomaszewska et al. 2023). Therefore, along with this evidence of aneuploidy and the large genetic distance of H031 from the other *U. humidicola* accessions (Jungmann et al. 2010), our results support the hypothesis proposed by Worthington et al. (2019) that H031 is a subspecies or a separate species.

Aneuploidy has been associated with unbalanced gametes and meiotic abnormalities (Soares et al. 2021), which, in turn, affect pollen production (Kaur and Singhal 2019; Zhang et al. 2013) and seed yield (Vleugels et al. 2019) in different species. In *Urochloa* spp., numerous studies have been carried out to understand the association between meiotic abnormalities and pollen fertility (da Cruz Baldissera et al. 2020; da Rocha et al. 2019; Souza et al. 2015). However, the extent to which such abnormalities influence seed yield is less well known. Improving seed yield, one of the major challenges faced by *U. humidicola* breeding programs, is an important goal that allows the establishment of cost-effective pastures, ensures the production of abundant progenies through intraspecific crosses and has economic implications for seed producers (Filho 1983; Ragalzi et al. 2021).

Previous studies evaluated seed production in hybrids of *U. humidicola*, all of which were generated from a cross between the sexual genotype H031 and pollen donor apomictic accessions from the EMBRAPA germplasm collection. The sexual hybrids were evaluated according to their meiotic abnormalities, and the authors observed a significant negative correlation between the percentage of meiotic abnormalities and the total production of pure seeds (Ragalzi et al. 2021). Similarly, the apomictic hybrids did not maintain the seed yield potential of their male parent (de Assis et al. 2016). Therefore, the aneuploidy observed in the H031 genotype may cause meiotic irregularities and a certain degree of genetic incompatibility with male apomicts, resulting in low seed production, poor germination, reduced seed viability and decreased yield.

In addition to meiotic stability, successful hybridization also requires parents with the same ploidy level, making it challenging to carry out intraspecific crosses in *U. humidicola*, since H031 is the only known hexaploid sexual accession and the apomictic cytotypes range from hexa-to nonaploids (Damasceno et al, 2023). The occurrence of natural sexual polyploid genotypes is also rare in other *Urochloa* species (Higgins et al. 2022; Hojsgaard and Hörandl 2015). Therefore, additional explorations of locations where the sexual genotype was documented may help us find valuable new material, with genetic variations that may be beneficial for enhancing the genetic diversity of *U. humidicola* and other *Urochloa* species (Valle and Pagliarini, 2009; Tomaszewska et al. 2023). In contrast to apomictic genotypes, which colonize large geographic regions through a process known as “geographical parthenogenesis”, sexual individuals typically remain limited to small areas (Higgins et al. 2022).

Despite the evidence of different genomic compositions in genotypes of *U. humidicola*, proposing a model for species evolution is much more difficult than in other *Urochloa* species because all accessions are polyploid and which diploid species should be considered ancestral is unclear (Tomaszewska et al. 2023). In this context, the use of techniques to visualize specific DNA sequences in chromosomes, such as cytomolecular analyses with genomic in situ hybridization (GISH) and molecular karyotyping through mapping by fluorescence in situ hybridization (FISH), integrated with sequencing data and bioinformatics tools, should be considered to provide insights into the origin and evolution of *U. humidicola* (Soltis et al, 2013; Damasceno et al, 2023).

Altogether, we reported an informative and high-quality genetic map and a candidate region for apomixis in the segmental allopolyploid *U. humidicola*. We hope that our results obtained using state-of-the-art algorithms for linkage analysis in polyploids will be relevant for the polyploid research community. We constructed haplotypes for all individuals, which have the potential to be applied in crop breeding, aiding in the identification of causal polymorphisms in a precise manner. We also detected preferential pairing in the apomictic accession H016 and its absence in the sexual accession H031. These results corroborate those of previous analyses and reinforce the need to better understand the origin of this sexual accession, since it is the only sexual genotype of *U. humidicola* and has been used as a female parent to generate hybrids in breeding programs. Moreover, 13 SNPs markers co-segregating to apomixis were identified and have potential for MAS. In summary, we provide valuable genetic and genomic resources that may facilitate the future molecular breeding of *U. humidicola* and other economically important tropical forage grasses.

## Supporting information

Moraes2023_SupplementaryMaterial

## Acknowledgments

We would like to acknowledge the Fundação de Amparo à Pesquisa do Estado de São Paulo (FAPESP), the Conselho Nacional de Desenvolvimento Científico e Tecnológico (CNPq), the Coordenação de Aperfeiçoamento de Pessoal de Nível Superior (CAPES), and the Empresa Brasileira de Pesquisa Agropecuária (Embrapa).

## Statements and declarations

### Data availability

The *U. humidicola* dataset presented in this study can be found at https://www.ncbi.nlm.nih.gov/, under the accession number PRJNA703438.

### Authors’ contributions

AAFG, CBV, APS and BBZV conceived the project and designed the experiments. ACLM, RCUF and BBZV performed the laboratory experiments. SCLB and CBV performed the field experiments. ACLM, MM, AA, LACL and MPF analyzed the data and interpreted the results. ACLM wrote the manuscript with major contributions from MM, RCFU and BBZV. All the authors have read and approved the manuscript.

## References

Aguilera PM, Galdeano F, Quarin CL, Amelio Ortiz JP and Espinoza F (2015). Inheritance of Aposporous Apomixis in Inter-specific Hybrids Derived from Sexual Paspalum plicatulum and Apomictic Paspalum guenoarum. Crop Science, 55(5), 1947–1956. 10.2135/cropsci2014.11.0770

Akiyama Y, Goel S, Conner JA, Hanna WW, Yamada-Akiyama H and Ozias-Akins P (2011) Evolution of the apomixis transmitting chromosome in pennisetum. BMC Evol Biol 11:289. 10.1186/1471-2148-11-289

Aono AH, Ferreira RCU, Moraes ADCL, Lara LADC, Pimenta RJG, Costa EA, Pinto LR, Landell MGA, Santos MF, Jank L, Barrios SCL, Valle CB, Chiari L, Garcia AAF, Kuroshu RM, Lorena AC, Gorjanc G and de Souza AP (2022) A joint learning approach for genomic prediction in polyploid grasses. Scientific reports, 12(1), 12499. 10.1038/s41598-022-16417-7

Assis G, dos Santos C, Silva P and Borges C (2014) Genetic divergence among Brachiaria humidicola (Rendle) schweick hybrids evaluated in the Western Brazilian Amazon. Crop Breed Appl Biotechnol 14:224–231. 10.1590/1984-70332014v14n4a35

Bennetzen JL, Schmutz J, Wang H et al (2012) Reference genome sequence of the model plant Setaria. Nat Biotechnol 30:555–561. 10.1038/nbt.2196

Bernini C and Marin-Morales MA (2001) Karyotype analysis in *Brachiaria* (Poaceae) species. Cytobios 104:157–171

Bhat JA, Yu D, Bohra A, Ganie SA and Varshney RK (2021) Features and applications of haplotypes in crop breeding. Commun Biol 4:1266. 10.1038/s42003-021-02782-y

Blischak PD, Kubatko LS and Wolfe AD (2018) SNP genotyping and parameter estimation in polyploids using low-coverage sequencing data. Bioinformatics 34:407–415. 10.1093/bioinformatics/btx587

Boldrini KR, Micheletti PL, Gallo PH, Mendes-Bonato AB, Pagliarini MS and Valle CB (2009) Origin of a polyploid accession of *Brachiaria* humidicola (Poaceae: Panicoideae: Paniceae). Genet Mol Res 8:888–895. 10.4238/vol8-3gmr617

Bomblies K (2023) Learning to tango with four (or more): The molecular basis of adaptation to polyploid meiosis. Plant Reprod 36:107–124. 10.1007/s00497-022-00448-1

Bomblies K, Jones G, Franklin C, Zickler D and Kleckner N (2016) The challenge of evolving stable polyploidy: Could an increase in “crossover interference distance” play a central role? Chromosoma 125:287–300. 10.1007/s00412-015-0571-4

Bourke PM, van Geest G, Voorrips RE, Jansen J, Kranenburg T, Shahin A, Visser RGF, Arens P, Smulders MJM and Maliepaard C (2018b) polymapR-linkage analysis and genetic map construction from F1 populations of outcrossing polyploids. Bioinformatics 34:3496–3502. 10.1093/bioinformatics/bty371

Bourke PM, Voorrips RE, Visser RGF and Maliepaard C (2018a) Tools for genetic studies in experimental populations of polyploids. Front Plant Sci 9:513. 10.3389/fpls.2018.00513

Choudhary A, Wright L, Ponce O, Chen J, Prashar A, Sanchez-Moran E, Luo Z and Compton L (2020) Varietal variation and chromosome behaviour during meiosis in solanum tuberosum. Heredity 125:212–226. 10.1038/s41437-020-0328-6

Conner JA, Goel S, Gunawan G et al (2008) Sequence analysis of bacterial artificial chromosome clones from the apospory-specific genomic region of pennisetum and cenchrus. Plant Physiol 147:1396–1411. 10.1104/pp.108.119081

Damasceno AG, Ferreira MTM, Soares IC, Barrios SCL, Do Valle CB, & Techio VH (2023). Physical mapping of ribosomal DNA sites and genome size in polyploid series of Urochloa humidicola (Rendle) Morrone & Zuloaga (Poaceae). Botany Letters, 1–10. 10.1080/23818107.2023.2192274

da Cruz Baldissera JN, Mendes ABD, Coan MMD, Mangolin CA, Do Valle CB and Pagliarini MS (2020) Selection based on meiotic behavior in urochloa decumbens hybrids from non-shattered seed. Trop Grassl Forrajes Trop 8:133–140. 10.17138/tgft(8)133-140

da Rocha MJ, Chiavegatto RB, Damasceno AG, Rocha LC, Souza Sobrinho F and Techio VH (2019) Comparative meiosis and cytogenomic analysis in euploid and aneuploid hybrids of urochloa P. beauv. Chromosome Res 27:333–344. 10.1007/s10577-019-09616-y

da Silva Pereira G, Mollinari M, Schumann MJ, Clough ME, Zeng ZB and Yencho GC (2021) The recombination landscape and multiple QTL mapping in a Solanum tuberosum cv. ‘Atlantic’-derived F(1) population. Heredity (Edinb) 126:817–830. 10.1038/s41437-021-00416-x

de Araujo Bitencourt, G., Chiari, L., & do Valle, C. B. (2012). Avaliação de híbridos por meio de marcadores RAPD e identificação do modo de reprodução pela anatomia de sacos embrionários em Brachiaria humidicola. Ensaios e Ciência C Biológicas Agrárias e da Saúde, 16(2).

de Assis GML, Beber PM, Clemencio RDM, Verzignassi J and do Valle CB (2016) Produção de sementes de genótipos de Brachiaria humidicola em Rio Branco, Acre. In: Congresso Brasileiro de Zootecnia, 26., 2016, Santa Maria, RS. Cinquenta anos de Zootecnia no Brasil: anais. Santa Maria, RS: SBZ, 2016..

de C Lara LA, Santos MF, Jank L, Chiari L, Vilela MM, Amadeu RR, Dos Santos JPR, Pereira GDS, Zeng ZB and Garcia AAF (2019) Genomic selection with allele dosage in panicum maximum jacq. G3 (Bethesda) 9:2463–2475. 10.1534/g3.118.200986

Deo TG, Ferreira RCU, Lara LAC, Moraes ACL, Alves-Pereira A, de Oliveira FA, Garcia AAF, Santos MF, Jank L and de Souza AP (2020) High-resolution linkage map with allele dosage allows the identification of regions governing complex traits and apospory in guinea grass (Megathyrsus maximus). Front Plant Sci 11:15. 10.3389/fpls.2020.00015

do Valle C, Bitencourt G, Chiari L, Resende R, Jank L and Arce A (2008) Identification of the mode of reproduction in Brachiaria humidicola hybrids. In International congress on sexual plant reproduction. Embrapa Recursos Genéticos e Biotecnologia, Brasília, pp 197

Doyle, J. J., & Doyle, J. L. (1987). A rapid DNA isolation procedure for small quantities of fresh leaf tissue. Phytochemical bulletin.

Elshire RJ, Glaubitz JC, Sun Q, Poland JA, Kawamoto K, Buckler ES and Mitchell SE (2011) A robust, simple genotyping-by-sequencing (GBS) approach for high diversity species. PLoS One 6:e19379. 10.1371/journal.pone.0019379

Ferreira RCU, da Costa Lima Moraes A, Chiari L, Simeão RM, Vigna BBZ and de Souza AP (2021) An overview of the genetics and genomics of the urochloa species most commonly used in pastures. Front Plant Sci 12:770461. 10.3389/fpls.2021.770461

Ferreira RCU, Lara LAC, Chiari L, Barrios SCL, do Valle CB, Valério JR, Torres FZV, Garcia AAF and de Souza AP (2019) Genetic mapping with allele dosage information in tetraploid urochloa decumbens (Stapf) R. D. Webster reveals insights into spittlebug (Notozulia entreriana Berg) resistance. Front Plant Sci 10:92. 10.3389/fpls.2019.00092

Filho MBD (1983) Limitações e potencial de Brachiaria humidicola para o trópico úmido brasileiro. Belém, Embrapa-Cpat

Galla G, Siena LA, Ortiz JPA, Baumlein H, Barcaccia G, Pessino SC, Bellucci M, and Pupilli F (2019). A portion of the apomixis locus of Paspalum simplex is microsyntenic with an unstable chromosome segment highly conserved among Poaceae. Scientific Reports, 9(1), 3271. 10.1038/s41598-019-39649-6

Gerard D, Ferrão LFV, Garcia AAF and Stephens M (2018) Genotyping polyploids from messy sequencing data. Genetics 210:789–807. 10.1534/genetics.118.301468

Glaubitz JC, Casstevens TM, Lu F, Harriman J, Elshire RJ, Sun Q, et al. (2014). TASSEL-GBS: a high capacity genotyping by sequencing analysis pipeline. PloS One 9, e90346. 10.1371/journal.pone.0090346

Goodstein DM, Shu S, Howson R, Neupane R, Hayes RD, Fazo J, Mitros T, Dirks W, Hellsten U, Putnam N and Rokhsar DS (2012) Phytozome: A comparative platform for green plant genomics. Nucleic Acids Res 40:D1178–D1186. 10.1093/nar/gkr944

Grandke F, Ranganathan S, van Bers N, de Haan JR and Metzler D (2017) PERGOLA: Fast and deterministic linkage mapping of polyploids. BMC Bioinform 18:12. 10.1186/s12859-016-1416-8

Gualtieri G, Conner JA, Morishige DT, Moore LD, Mullet JE and Ozias-Akins P (2006) A segment of the apospory-specific genomic region is highly microsyntenic not only between the apomicts pennisetum squamulatum and buffelgrass, but also with a rice chromosome 11 centromeric-proximal genomic region. Plant Physiol 140:963–971. 10.1104/pp.105.073809

Hackett CA, Boskamp B, Vogogias A, Preedy KF and Milne I (2017) TetraploidSNPMap: Software for linkage analysis and QTL mapping in autotetraploid populations using SNP dosage data. J Hered 108:438–442. 10.1093/jhered/esx022

Henderson IR (2012) Control of meiotic recombination frequency in plant genomes. Current opinion in plant biology, 15(5), 556–561. 10.1016/j.pbi.2012.09.002

Higgins J, Tomaszewska P, Pellny TK, Castiblanco V, Arango J, Tohme J, Schwarzacher T, Mitchell RA, Heslop-Harrison JS and De Vega JJ (2022) Diverged subpopulations in tropical urochloa (*Brachiaria*) forage species indicate a role for facultative apomixis and varying ploidy in their population structure and evolution. Ann Bot 130:657–669. 10.1093/aob/mcac115

Hojsgaard D and Hörandl E (2015) Apomixis as a facilitator of range expansion and diversification in plants. In: Pontarotti P (ed) Evolutionary biology: Biodiversification from genotype to phenotype. Springer International Publishing, Cham, pp 305–327

Ishigaki G, Gondo T, Ebina M, Suenaga K and Akashi R (2010) Estimation of genome size in Brachiaria species. Grassland Science, 56(4), 240–242. 10.1111/j.1744-697X.2010.00200.x

Jensen SE, Charles JR, Muleta K et al (2020) A sorghum practical haplotype graph facilitates genome-wide imputation and cost-effective genomic prediction. Plant Genome 13:e20009. 10.1002/tpg2.20009

Jungmann L, Vigna BB, Boldrini KR, Sousa AC, do Valle CB, Resende RM, Pagliarini MS, Zucchi MI and de Souza AP (2010) Genetic diversity and population structure analysis of the tropical pasture grass Brachiaria humidicola based on microsatellites, cytogenetics, morphological traits, and geographical origin. Genome 53:698–709. 10.1139/g10-055

Kamiri M, Stift M, Costantino G, Dambier D, Kabbage T, Ollitrault P and Froelicher Y (2018) Preferential homologous chromosome pairing in a tetraploid intergeneric somatic hybrid (*Citrus reticulata* + *Poncirus trifoliata*) revealed by molecular marker inheritance. Front Plant Sci 9:1557. 10.3389/fpls.2018.01557

Kaur D and Singhal VK (2019) Meiotic abnormalities affect genetic constitution and pollen viability in dicots from Indian cold deserts. BMC Plant Biol 19:10. 10.1186/s12870-018-1596-7

Kaushal P, Dwivedi KK, Radhakrishna A, Srivastava MK, Kumar V, Roy AK and Malaviya DR (2019) Partitioning apomixis components to understand and utilize gametophytic apomixis. Front Plant Sci 10:256. 10.3389/fpls.2019.00256

Langmead, B., Trapnell, C., Pop, M., Salzberg, S. L. (2009). Ultrafast and memory-efficient alignment of short DNA sequences to the human genome. Genome Biol. 10 (3), R25. 10.1186/gb-2009-10-3-r25

Leach LJ, Wang L, Kearsey MJ and Luo Z (2010) Multilocus tetrasomic linkage analysis using hidden markov chain model. Proc Natl Acad Sci U S A 107:4270–4274. 10.1073/pnas.0908477107

Lenormand T, Engelstädter J, Johnston SE, Wijnker E and Haag CR (2016) Evolutionary mysteries in meiosis. Philos Trans R Soc Lond B Biol Sci 371:20160001. 10.1098/rstb.2016.0001

Liao Y, Voorrips RE, Bourke PM, Tumino G, Arens P, Visser RGF, Smulders MJM and Maliepaard C (2021) Using probabilistic genotypes in linkage analysis of polyploids. Theor Appl Genet 134:2443–2457. 10.1007/s00122-021-03834-x

Majidian S, Kahaei MH and de Ridder D (2020) Hap10: Reconstructing accurate and long polyploid haplotypes using linked reads. BMC Bioinform 21:253. 10.1186/s12859-020-03584-5

Martins FB, Moraes ACL, Aono AH, Ferreira RCU, Chiari L, Simeão RM, Barrios SCL, Santos MF, Jank L, do Valle CB, Vigna BBZ and de Souza AP (2021) A semi-automated SNP-based approach for contaminant identification in biparental polyploid populations of tropical forage grasses. Front Plant Sci 12:737919. 10.3389/fpls.2021.737919

Mason AS and Wendel JF (2020) Homoeologous exchanges, segmental allopolyploidy, and polyploid genome evolution. Front Genet 11:1014. 10.3389/fgene.2020.01014

Matias FI, Alves FC, Meireles KGX, Barrios SCL, do Valle CB, Endelman JB and Fritsche-Neto R (2019b) On the accuracy of genomic prediction models considering multi-trait and allele dosage in urochloa spp. interspecific tetraploid hybrids. Mol Breed 39:100. 10.1007/s11032-019-1002-7

Matias FI, Vidotti MS, Meireles KGX, Barrios SCL, do Valle CB, Carley CAS and Fritsche-Neto R (2019a) Association mapping considering allele dosage: An example of forage traits in an interspecific segmental allotetraploid urochloa spp. Panel. Crop Sci 59:2062–2076. 10.2135/cropsci2019.03.0185

Mayer M, Hölker AC, González-Segovia E, Bauer E, Presterl T, Ouzunova M, Melchinger AE and Schön C-C (2020) Discovery of beneficial haplotypes for complex traits in maize landraces. Nat Commun 11:4954. 10.1038/s41467-020-18683-3

Mollinari M and Garcia AAF (2019) Linkage analysis and haplotype phasing in experimental autopolyploid populations with high ploidy level using hidden markov models. G3 (Bethesda) 9:3297–3314. 10.1534/g3.119.400378

Mollinari M, Olukolu BA, Pereira GDS, Khan A, Gemenet D, Yencho GC and Zeng ZB (2020) Unraveling the hexaploid sweetpotato inheritance using ultra-dense multilocus mapping. G3 (Bethesda) 10:281-292. 10.1534/g3.119.400620

Moreau CS (2014) A practical guide to DNA extraction, PCR, and gene-based DNA sequencing in insects. Halteres 5:32–42

Okada M, Lanzatella C, Saha MC, Bouton J, Wu R and Tobias CM (2010) Complete switchgrass genetic maps reveal subgenome collinearity, preferential pairing and multilocus interactions. Genetics 185:745–760. 10.1534/genetics.110.113910

Oloka BM, da Silva Pereira G, Amankwaah VA, Mollinari M, Pecota KV, Yada B, Olukolu BA, Zeng ZB and Craig Yencho G (2021) Discovery of a major QTL for root-knot nematode (Meloidogyne incognita) resistance in cultivated sweetpotato (Ipomoea batatas). Theor Appl Genet 134:1945–1955. 10.1007/s00122-021-03797-z

Ortiz JPA, Pupilli F, Acuña CA, Leblanc O and Pessino SC (2020) How to become an apomixis model: The multifaceted case of paspalum. Genes (Basel) 11:974. 10.3390/genes11090974

Ozias-Akins P, Akiyama Y and Hanna WW (2003) Molecular characterization of the genomic region linked with apomixis in Pennisetum/Cenchrus. Functional & integrative genomics, 3, 94–104. 10.1007/s10142-003-0084-8

Ozias-Akins P and van Dijk PJ (2007) Mendelian genetics of apomixis in plants. Annu Rev Genet 41:509–537. 10.1146/annurev.genet.40.110405.090511

Palumbo F, Draga S, Vannozzi A, Lucchin M and Barcaccia G (2022) Trends in apomixis research: The 10 most cited research articles published in the pregenomic and genomic eras. Front Plant Sci 13:878074. 10.3389/fpls.2022.878074

Patel RK, & Jain M (2012) NGS QC Toolkit: a toolkit for quality control of next generation sequencing data. PloS one, 7(2), e30619. 10.1371/journal.pone.0030619

Penteado MdO, dos Santos A, Rodrigues IF, do Valle C, Seixas MAC and Esteves A (2000) Determinação de ploidia e avaliação da quantidade de DNA total em diferentes espécies do gênero Brachiaria. Campo Grande, Embrapa Gado de Corte

Pereira GS, Garcia AAF and Margarido GRA (2018b) A fully automated pipeline for quantitative genotype calling from next generation sequencing data in autopolyploids. BMC Bioinform 19:398. 10.1186/s12859-018-2433-6

Pereira JF, Azevedo ALS, Pessoa-Filho M et al (2018a) Research priorities for next-generation breeding of tropical forages in Brazil. Crop Breed Appl Biotechnol 18:314–319. 10.1590/1984-70332018v18n3n46

Pessino SC, Evans C, Ortiz JPA, Armstead I, Valle CBD and Hayward MD (1998) A genetic Map of the apospory-region in brachiaria hybrids: identification of two markers closely associated with the trait. Hereditas, 128(2), 153–158. 10.1111/j.1601-5223.1998.00153.x

Pessoa-Filho M, Martins AM and Ferreira ME (2017) Molecular dating of phylogenetic divergence between urochloa species based on complete chloroplast genomes. BMC Genom 18:516. 10.1186/s12864-017-3904-2

Pessoa-Filho M, Sobrinho FS, Fragoso RR, Silva Junior OB and Ferreira ME (2019) A phased diploid genome assembly for the forage grass Urochloa ruziziensis based on single-molecule real-time sequencing. In International Plant and Animal Genome Conference XXVII, 2019. San Diego

Poland JA, Brown PJ, Sorrells ME and Jannink JL (2012) Development of high-density genetic maps for barley and wheat using a novel two-enzyme genotyping-by-sequencing approach. PLoS One 7:e32253. 10.1371/journal.pone.0032253

Preedy, KF, & Hackett, CA (2016) A rapid marker ordering approach for high-density genetic linkage maps in experimental autotetraploid populations using multidimensional scaling. Theoretical and Applied Genetics, 129, 2117–2132. 10.1007/s00122-016-2761-8

Ragalzi CDM, Mendes ABD, Simeão RM, Verzignassi JR, Valle CBD and Machado MDFPDS (2021) Microsporogenesis associated with seed yield in urochloa sexual polyploid hybrids. Crop Breed Appl Biotechnol 21:e37652148. 10.1590/1984-70332021v21n4a57

Schmitz A, & Riesner D (2006) Purification of nucleic acids by selective precipitation with polyethylene glycol 6000. Analytical biochemistry, 354(2), 311–313. 10.1016/j.ab.2006.03.014

Serang O, Mollinari M and Garcia AA (2012) Efficient exact maximum a posteriori computation for bayesian SNP genotyping in polyploids. PLoS One 7:e30906. 10.1371/journal.pone.0030906

Simeão RM, Resende MDV, Alves RS, Pessoa-Filho M, Azevedo ALS, Jones CS, Pereira JF and Machado JC (2021) Genomic selection in tropical forage grasses: Current status and future applications. Front Plant Sci 12:665195. 10.3389/fpls.2021.665195

Soares NR, Mollinari M, Oliveira GK, Pereira GS and Vieira MLC (2021) Meiosis in polyploids and implications for genetic mapping: A review. Genes (Basel) 12:1517. 10.3390/genes12101517

Soltis DE, Gitzendanner MA, Stull G, Chester M, Chanderbali A, Chamala S, Jordon-Thaden I, Soltis PS, Schnable PS and Barbazuk WB (2013) The potential of genomics in plant systematics. Taxon, 62(5), 886–898. 10.12705/625.13

Souza VF, Pagliarini MS, Valle CB, Bione NC, Menon MU and Mendes-Bonato AB (2015) Meiotic behavior of *Brachiaria decumbens* hybrids. Genet Mol Res 14:12855–12865. 10.4238/2015.October.21.5

Tomaszewska P, Vorontsova MS, Renvoize SA, Ficinski SZ, Tohme J, Schwarzacher T, Castiblanco V, de Vega JJ, Mitchell RAC and Heslop-Harrison JSP (2023) Complex polyploid and hybrid species in an apomictic and sexual tropical forage grass group: Genomic composition and evolution in urochloa (*Brachiaria*) species. Ann Bot 131:87–108. 10.1093/aob/mcab147

Triviño NJ, Perez JG, Recio ME, Ebina M, Yamanaka N, Tsuruta S-I, Ishitani M and Worthington M (2017) Genetic diversity and population structure of B*rachiaria* species and breeding populations. Crop Sci 57:2633–2644. 10.2135/cropsci2017.01.0045

Valle CD and Pagliarini MS (2009) Biology, cytogenetics, and breeding of Brachiaria. Genetic resources, chromosome engineering, and crop improvement, 5

Vigna BB, Santos JC, Jungmann L, do Valle CB, Mollinari M, Pastina MM, Pagliarini MS, Garcia AA and Souza AP (2016) Evidence of allopolyploidy in urochloa humidicola based on cytological analysis and genetic linkage mapping. PLoS One 11:e0153764. 10.1371/journal.pone.0153764

Vleugels T, Laere KV, Roldán-Ruiz I and Cnops G (2019) Seed yield in red clover is associated with meiotic abnormalities and in tetraploid genotypes also with self-compatibility. Euphytica 215:79. 10.1007/s10681-019-2405-6

Worthington M, Ebina M, Yamanaka N et al (2019) Translocation of a parthenogenesis gene candidate to an alternate carrier chromosome in apomictic *Brachiaria humidicola*. BMC Genom 20:41. 10.1186/s12864-018-5392-4

Worthington M, Heffelfinger C, Bernal D, Quintero C, Zapata YP, Perez JG, De Vega J, Miles J, Dellaporta S and Tohme J (2016) A parthenogenesis gene candidate and evidence for segmental allopolyploidy in apomictic B*rachiaria decumbens*. Genetics 203:1117–1132. 10.1534/genetics.116.190314

Worthington M, Perez JG, Mussurova S et al (2021) A new genome allows the identification of genes associated with natural variation in aluminium tolerance in *Brachiaria* grasses. J Exp Bot 72:302–319. 10.1093/jxb/eraa469

Wu KK, Burnquist W, Sorrells ME, Tew TL, Moore PH and Tanksley SD (1992) The detection and estimation of linkage in polyploids using single-dose restriction fragments. Theoretical and Applied Genetics, 83, 294–300. 10.1007/BF00224274

Xiong J, Hu F, Ren J, Huang Y, Liu C and Wang K (2023) Synthetic apomixis: The beginning of a new era. Curr Opin Biotechnol 79:102877. 10.1016/j.copbio.2022.102877

Young BA, Sherwood RT and Bashaw EC (1979) Cleared-pistil and thick-sectioning techniques for detecting aposporous apomixis in grasses. Can J Bot 57:1668–1672. 10.1139/b79-204

Zhang G, Liu X, Quan Z et al (2012) Genome sequence of foxtail millet (Setaria italica) provides insights into grass evolution and biofuel potential. Nat Biotechnol 30:549–554. 10.1038/nbt.2195

Zhang H, Bian Y, Gou X et al (2013) Persistent whole-chromosome aneuploidy is generally associated with nascent allohexaploid wheat. Proc Natl Acad Sci U S A 110:3447–3452. 10.1073/pnas.1300153110

Zheng C, Amadeu RR, Munoz PR and Endelman JB (2021) Haplotype reconstruction in connected tetraploid F1 populations. Genetics 219:iyab106. 10.1093/genetics/iyab106

Zielinski M-L and Mittelsten Scheid O (2012) Meiosis in polyploid plants. In: Soltis PS, Soltis DE (ed) Polyploidy and genome evolution. Springer Berlin Heidelberg, Berlin, Heidelberg, pp 33–55

Zorzatto C, Chiari L, De Araújo Bitencourt G, Do Valle CB, De Campos Leguizamón GO, Schuster I and Pagliarini MS (2010) Identification of a molecular marker linked to apomixis in Brachiaria humidicola (Poaceae). Plant Breed 129:734–736. 10.1111/j.1439-0523.2010.01763.x

