## Supplementary material for "Advances in genomic characterization of *Urochloa humidicola*: exploring polyploid inheritance and apomixis": Moraes2023_SupplementaryMaterial

**Supplementary information**

**Supplementary figures**

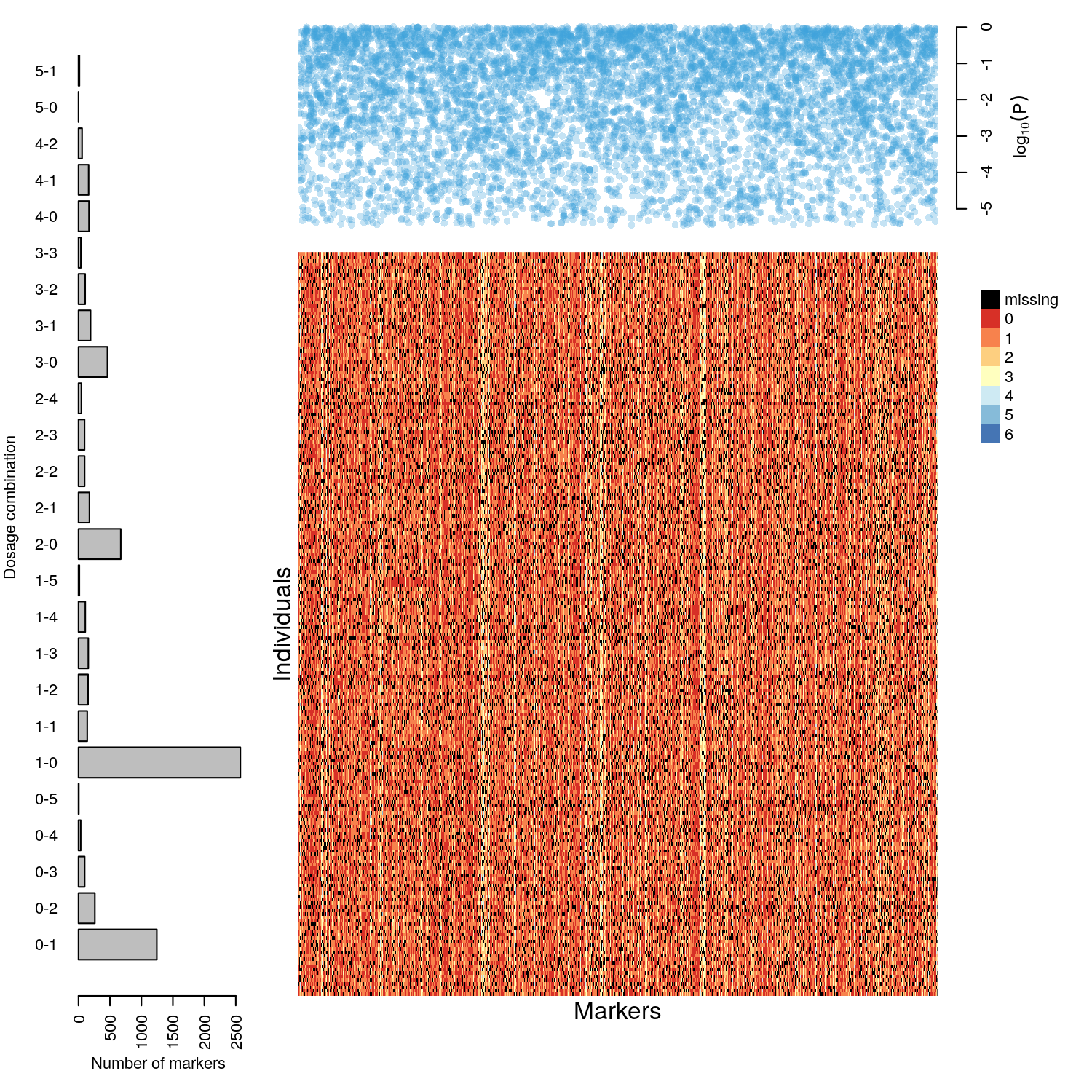

**Supplementary Fig. 1** Distribution of the 7,067 high-quality SNPs with allele dosage information retained for linkage analyses in *U. humidicola* hybrids. Of these SNPs, 54.2% were classified as simplex, 2.4% as double-simplex, and 43.5% as higher-dosage markers (multiplex)

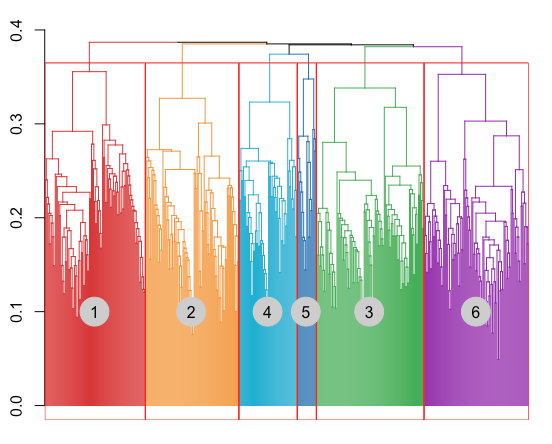

**Supplementary Fig. 2** Dendrogram showing six LGs obtained using the UPGMA clustering method. Numbers in the dendrogram indicate the six LGs of *U. humidicola*

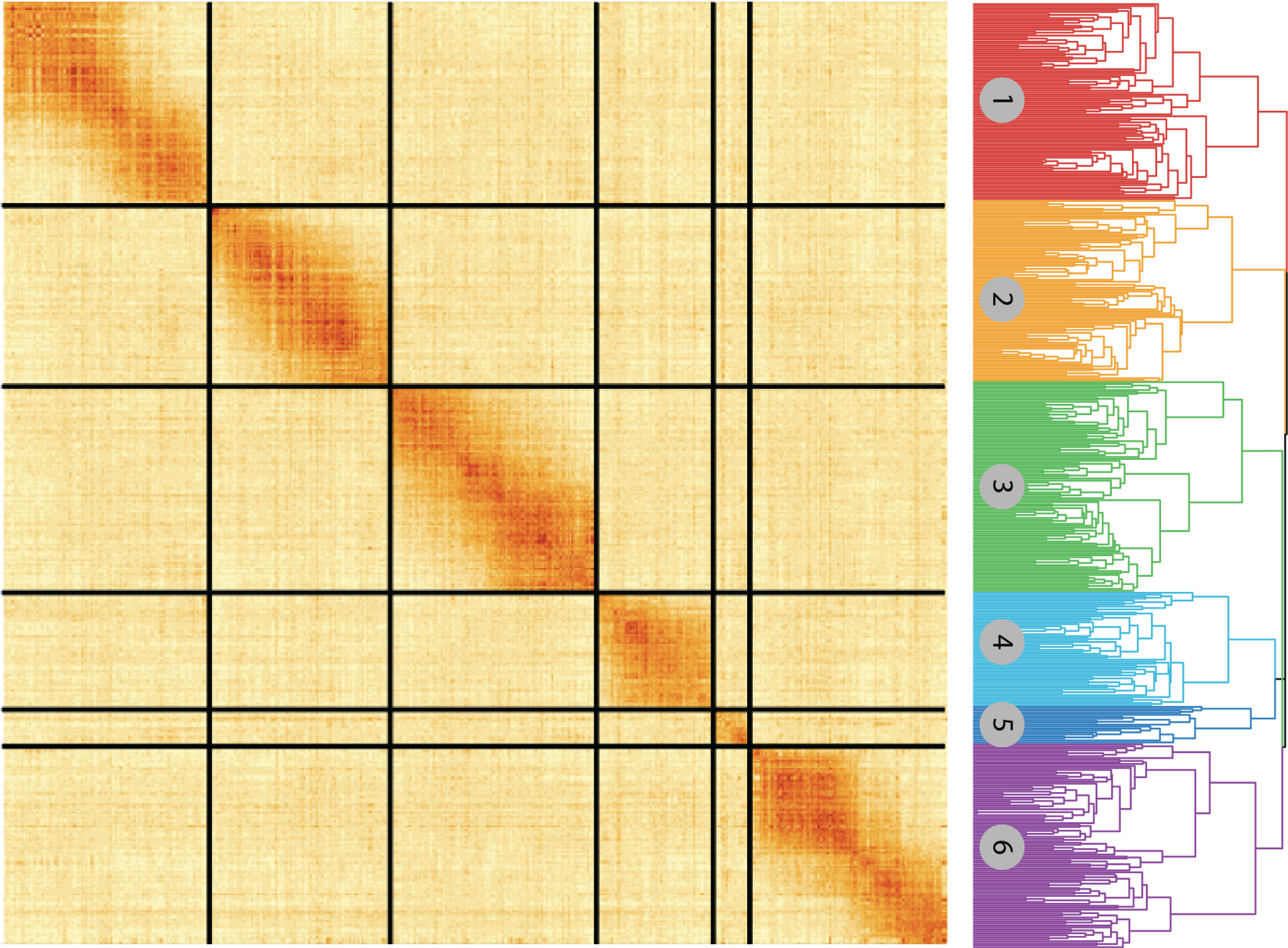

**Supplementary Fig. 3** Aggregated recombination fraction between 4,802 SNPs averaged into a grid of 10x10 for enhanced visibility of linkage blocks. The matrix was clustered using the unweighted pair group method with arithmetic mean (UPGMA) algorithm to generate SNP clusters corresponding to the linkage groups. For each linkage group, we applied the multidimensional scaling algorithm. The six submatrices along the diagonal represent the ordered linkage groups of *U. humidicola*

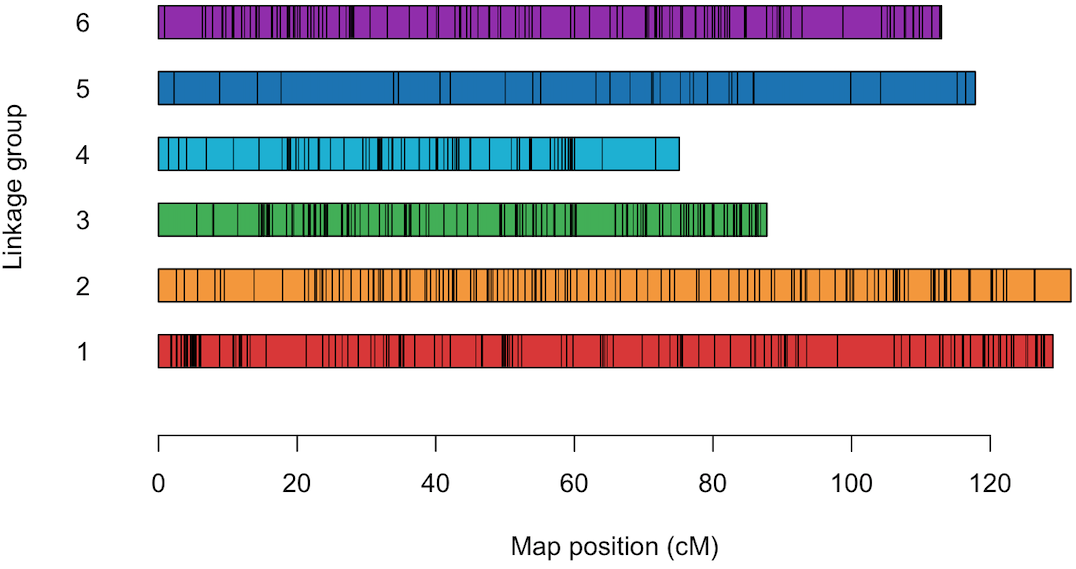

### **Supplementary Fig. 4** The genetic linkage map of *U. humidicol*a constructed using SNP markers with allele dosage information. Black bars represent SNP markers. The x-axis represents the genetic position in centimorgans (cM), while the y-axis represents the linkage group

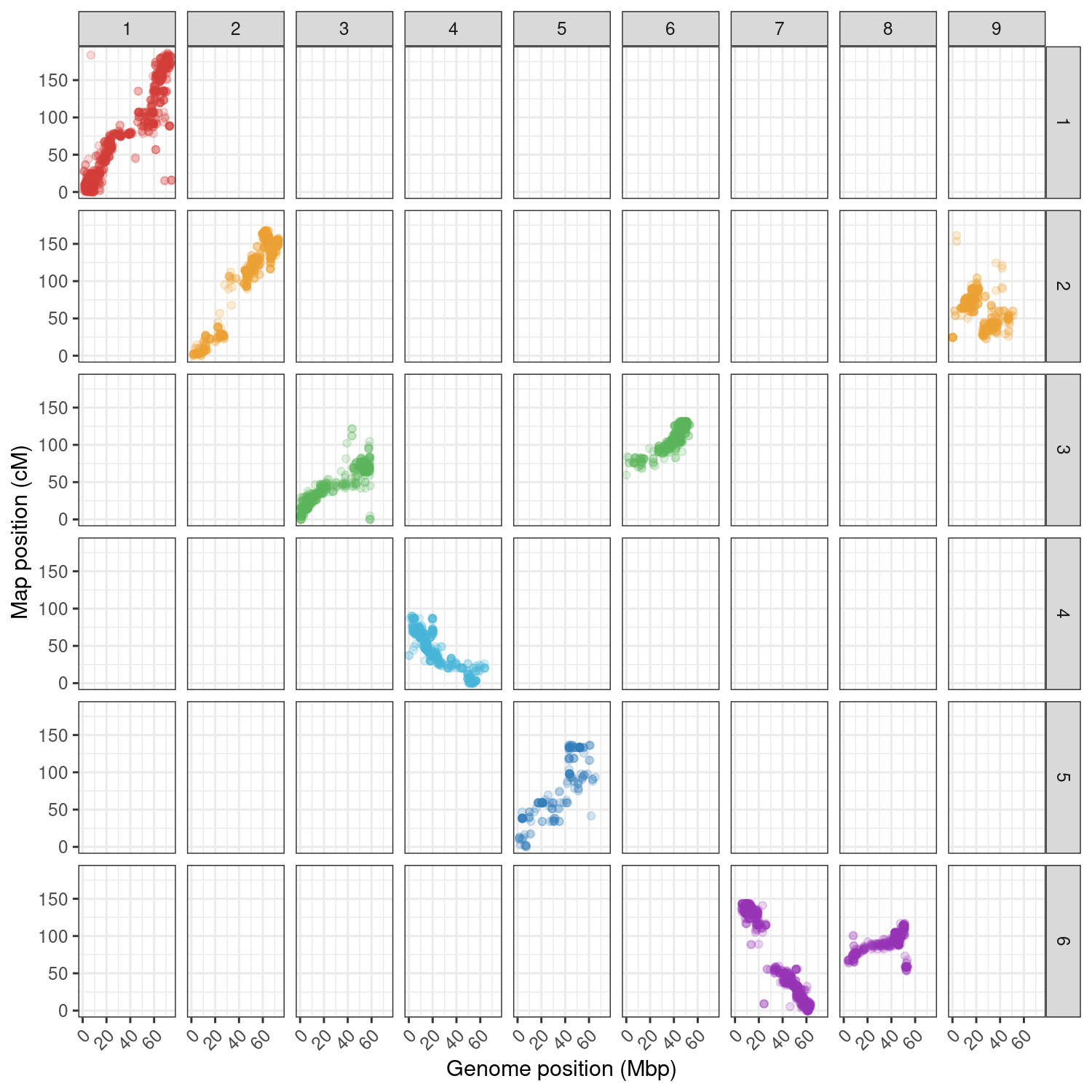

**Supplementary Fig. 5** Genetic distance in the six linkage groups of *U. humidicola* (map position in cM) versus physical distance in the nine chromosome of *U. ruziziensis* reference genome

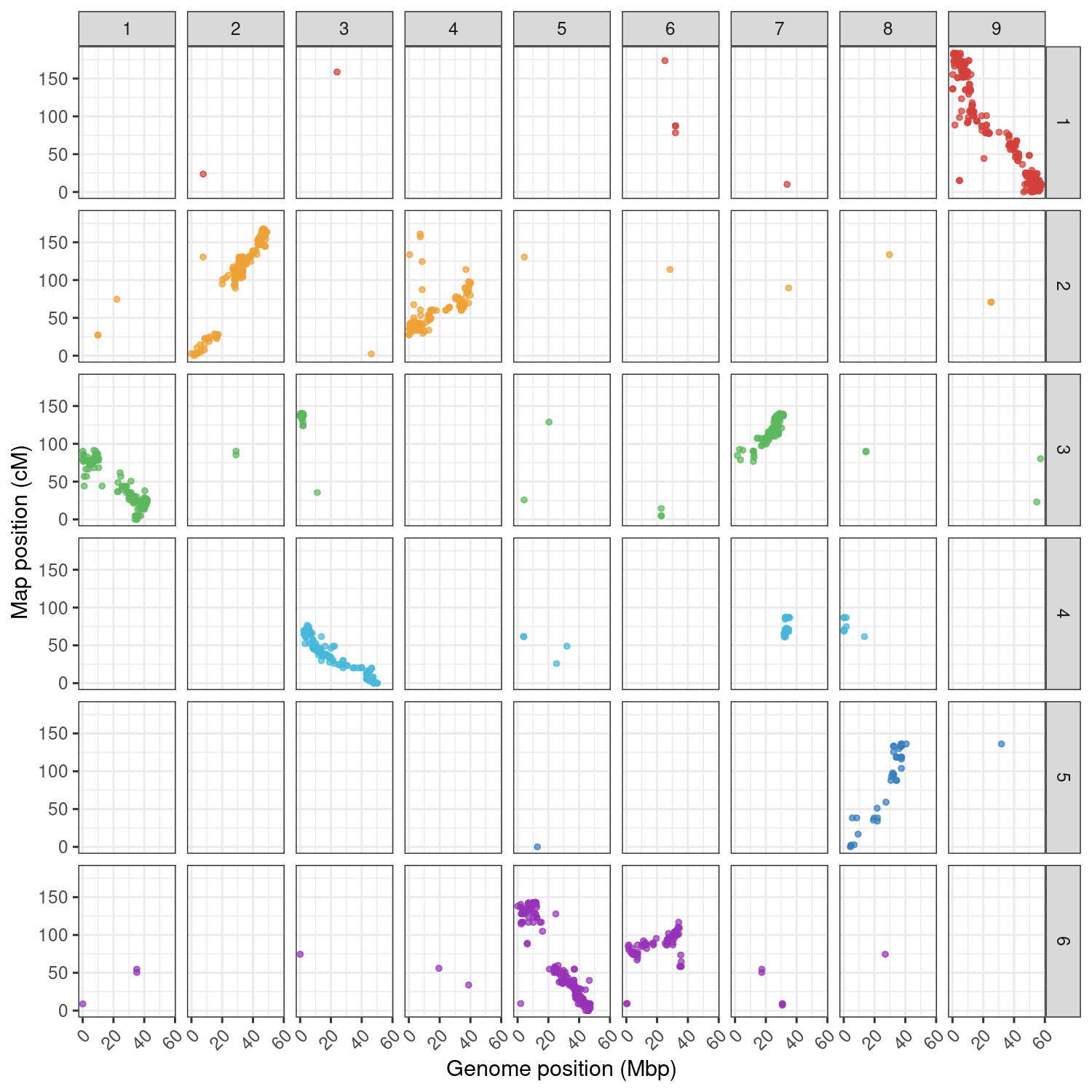

**Supplementary Fig. 6** Genetic distance in the six linkage groups of *U. humidicola* (map position in cM) versus physical distance in the nine chromosomes of *S. italica* genome

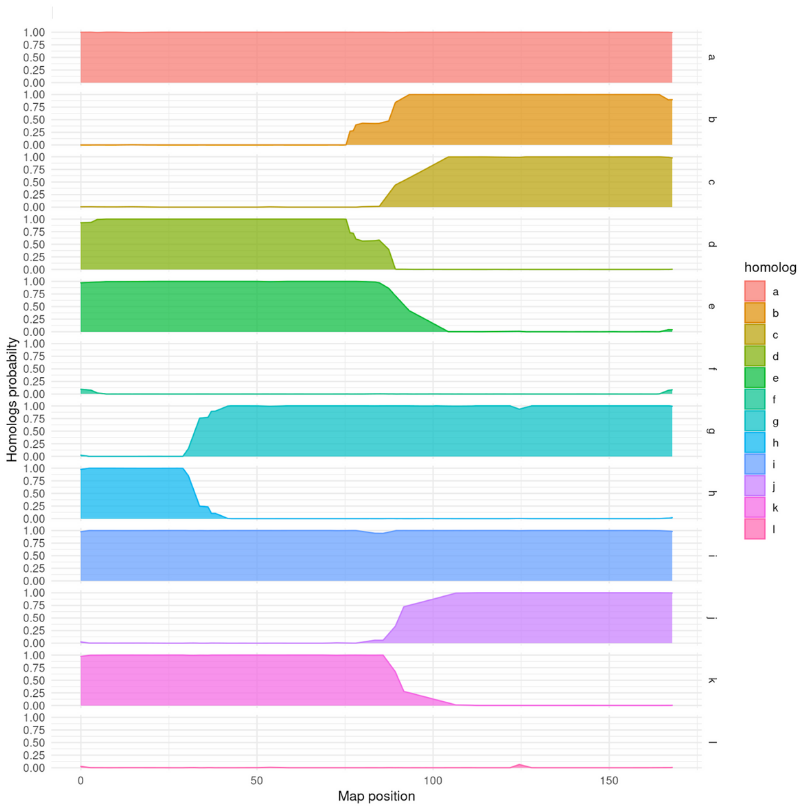

**Supplementary Fig. 7** Example of haplotype reconstruction and distribution of meiotic configurations for individual Bh1, linkage group 2. The probability profiles for 12 homologs indicate the segments inherited from the parents "H016" (homologs "a" to "f") and "H031" ("g" to "l")

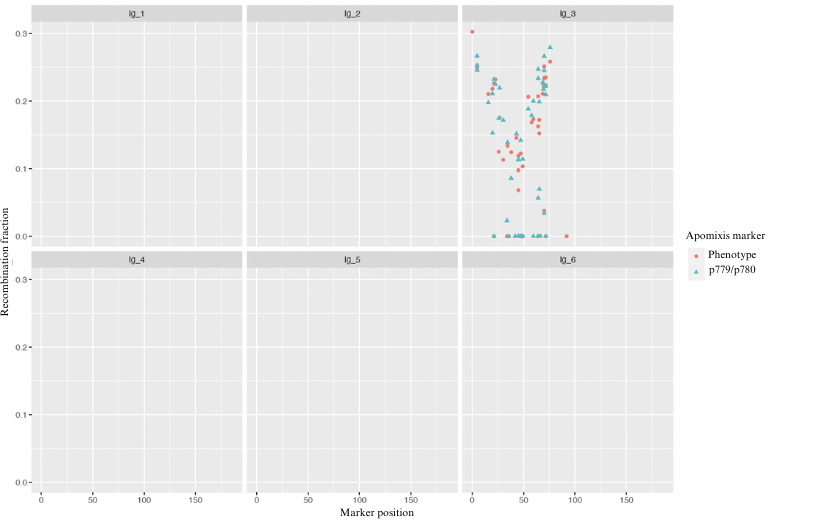

**Supplementary Fig. 8** The ASGR-specific marker p779/p780 assigned to position 53.73 cM of LG3 and the apomixis phenotype assigned to position 47.16 of the same group

**Supplementary Table S1** Genotype scores of the ASGR–BBML-specific marker p779/p780 evaluated in the parents and hybrids of the *U. humidicola* population

| **Individual** | **Phenotype*** | **Expected score based on the phenotype**** | **Observed score using the p779/780 marker** |
| --- | --- | --- | --- |
| Bh1 | apomictic | 1 | 1 |
| Bh2 | sexual | 0 | 0 |
| Bh3 | apomictic | 1 | 1 |
| Bh4 | apomictic | 1 | 0 |
| Bh5 | apomictic | 1 | 1 |
| Bh6 | sexual | 0 | 1 |
| Bh7 | apomictic | 1 | 1 |
| Bh8 | apomictic | 1 | 1 |
| Bh9 | apomictic | 1 | 1 |
| Bh11 | sexual | 0 | 0 |
| Bh12 | apomictic | 1 | 1 |
| Bh14 | sexual | 0 | 0 |
| Bh16 | sexual | 0 | 0 |
| Bh18 | apomictic | 1 | 1 |
| Bh20 | sexual | 0 | 1 |
| Bh21 | apomictic | 1 | 1 |
| Bh23 | sexual | 0 | 0 |
| Bh24 | sexual | 0 | 1 |
| Bh25 | apomictic | 1 | 1 |
| Bh26 | sexual | 0 | 0 |
| Bh27 | sexual | 0 | 0 |
| Bh28 | apomictic | 1 | 0 |
| Bh29 | apomictic | 1 | 1 |
| Bh30 | apomictic | 1 | 1 |
| Bh31 | apomictic | 1 | 1 |
| Bh32 | sexual | 0 | 0 |
| Bh33 | sexual | 0 | 0 |
| Bh34 | sexual | 0 | 0 |
| Bh35 | sexual | 0 | 0 |
| Bh36 | apomictic | 1 | 1 |
| Bh37 | apomictic | 1 | 1 |
| Bh39 | sexual | 0 | 0 |
| Bh41 | sexual | 0 | 0 |
| Bh43 | apomictic | 1 | 1 |
| Bh45 | sexual | 0 | 0 |
| Bh47 | apomictic | 1 | 1 |
| Bh54 | apomictic | 1 | 1 |
| Bh60 | sexual | 0 | 0 |
| Bh61 | apomictic | 1 | 1 |
| Bh63 | sexual | 0 | 1 |
| Bh64 | sexual | 0 | 0 |
| Bh65 | apomictic | 1 | 1 |
| Bh68 | apomictic | 1 | 1 |
| Bh69 | sexual | 0 | 0 |
| Bh72 | sexual | 0 | 0 |
| Bh73 | sexual | 0 | 0 |
| Bh76 | sexual | 0 | 0 |
| Bh77 | sexual | 0 | 0 |
| Bh78 | sexual | 0 | 0 |
| Bh80 | apomictic | 1 | 1 |
| Bh81 | sexual | 0 | 0 |
| Bh83 | apomictic | 1 | 1 |
| Bh84 | sexual | 0 | 1 |
| Bh85 | apomictic | 1 | 1 |
| Bh87 | sexual | 0 | 0 |
| Bh88 | apomictic | 1 | 1 |
| Bh92 | apomictic | 1 | 1 |
| Bh93 | sexual | 0 | 0 |
| Bh94 | sexual | 0 | 1 |
| Bh95 | sexual | 0 | 1 |
| Bh96 | apomictic | 1 | 1 |
| Bh97 | apomictic | 1 | 1 |
| Bh98 | apomictic | 1 | 1 |
| Bh99 | apomictic | 1 | 1 |
| Bh100 | apomictic | 1 | 1 |
| Bh101 | apomictic | 1 | 1 |
| Bh102 | apomictic | 1 | 1 |
| Bh103 | sexual | 0 | 0 |
| Bh104 | sexual | 0 | 1 |
| Bh105 | sexual | 0 | 0 |
| Bh106 | apomictic | 1 | 1 |
| Bh108 | sexual | 0 | 0 |
| Bh109 | apomictic | 1 | 1 |
| Bh110 | apomictic | 1 | 1 |
| Bh111 | apomictic | 1 | 1 |
| Bh112 | sexual | 0 | 1 |
| Bh113 | apomictic | 1 | 1 |
| Bh114 | apomictic | 1 | 1 |
| Bh115 | apomictic | 1 | 1 |
| Bh117 | sexual | 0 | 0 |
| Bh119 | sexual | 0 | 0 |
| Bh120 | apomictic | 1 | 1 |
| Bh121 | sexual | 0 | 0 |
| Bh122 | apomictic | 1 | 0 |
| Bh123 | sexual | 0 | 0 |
| Bh124 | apomictic | 1 | 1 |
| Bh125 | apomictic | 1 | 0 |
| Bh126 | sexual | 0 | 0 |
| Bh127 | sexual | 0 | 1 |
| Bh128 | sexual | 0 | 1 |
| Bh129 | sexual | 0 | 0 |
| Bh130 | sexual | 0 | 0 |
| Bh131 | sexual | 0 | 0 |
| Bh132 | sexual | 0 | 0 |
| Bh133 | sexual | 0 | 0 |
| Bh134 | sexual | 0 | 0 |
| Bh135 | sexual | 0 | 0 |
| Bh136 | apomictic | 1 | 1 |
| Bh137 | sexual | 0 | 0 |
| Bh138 | sexual | 0 | 1 |
| Bh140 | sexual | 0 | 0 |
| Bh141 | sexual | 0 | 0 |
| Bh142 | sexual | 0 | 1 |
| Bh143 | apomictic | 1 | 1 |
| Bh144 | sexual | 0 | 0 |
| Bh145 | apomictic | 1 | 1 |
| Bh146 | apomictic | 1 | 1 |
| Bh147 | sexual | 0 | 0 |
| Bh148 | sexual | 0 | 0 |
| Bh149 | apomictic | 1 | 1 |
| Bh150 | apomictic | 1 | 1 |
| Bh151 | sexual | 0 | 0 |
| Bh152 | apomictic | 1 | 1 |
| Bh153 | apomictic | 1 | 1 |
| Bh154 | apomictic | 1 | 1 |
| Bh155 | apomictic | 1 | 0 |
| Bh156 | apomictic | 1 | 0 |
| Bh157 | sexual | 0 | 0 |
| Bh158 | apomictic | 1 | 1 |
| Bh159 | apomictic | 1 | 1 |
| Bh160 | sexual | 0 | 0 |
| Bh161 | sexual | 0 | 0 |
| Bh162 | sexual | 0 | 0 |
| Bh163 | apomictic | 1 | 1 |
| Bh164 | sexual | 0 | 1 |
| Bh165 | sexual | 0 | 0 |
| Bh166 | sexual | 0 | 0 |
| Bh167 | sexual | 0 | 0 |
| Bh168 | sexual | 0 | 0 |
| Bh169 | sexual | 0 | 0 |
| Bh170 | apomictic | 1 | 1 |
| Bh172 | sexual | 0 | 0 |
| Bh175 | sexual | 0 | 1 |
| Bh176 | apomictic | 1 | 1 |
| Bh177 | sexual | 0 | 1 |
| Bh178 | apomictic | 1 | 1 |
| Bh179 | sexual | 0 | 0 |
| Bh180 | sexual | 0 | 1 |
| Bh181 | sexual | 0 | 0 |
| Bh182 | sexual | 0 | 0 |
| Bh183 | apomictic | 1 | 1 |
| Bh184 | sexual | 0 | 0 |
| Bh185 | sexual | 0 | 0 |
| Bh187 | sexual | 0 | 1 |
| Bh188 | apomictic | 1 | 1 |
| Bh189 | sexual | 0 | 0 |
| Bh190 | apomictic | 1 | 1 |
| Bh191 | apomictic | 1 | 0 |
| Bh192 | sexual | 0 | 0 |
| Bh193 | sexual | 0 | 0 |
| Bh196 | sexual | 0 | 0 |
| Bh199 | apomictic | 1 | 1 |
| Bh200 | apomictic | 1 | 1 |
| Bh202 | apomictic | 1 | 1 |
| Bh203 | sexual | 0 | 0 |
| Bh204 | sexual | 0 | 0 |
| Bh205 | apomictic | 1 | 0 |
| Bh206 | apomictic | 1 | 1 |
| Bh207 | apomictic | 1 | 1 |
| Bh208 | sexual | 0 | 0 |
| Bh209 | sexual | 0 | 1 |
| Bh210 | apomictic | 1 | 1 |
| Bh211 | sexual | 0 | 0 |
| Bh212 | sexual | 0 | 0 |
| Bh213 | sexual | 0 | 1 |
| Bh214 | sexual | 0 | 0 |
| Bh215 | apomictic | 1 | 1 |
| Bh216 | sexual | 0 | 0 |
| Bh217 | apomictic | 1 | 1 |
| Bh218 | sexual | 0 | 0 |
| Bh219 | sexual | 0 | 0 |
| Bh220 | sexual | 0 | 0 |
| Bh221 | sexual | 0 | 0 |
| Bh222 | sexual | 0 | 1 |
| Bh223 | sexual | 0 | 0 |
| Bh224 | sexual | 0 | 0 |
| Bh225 | sexual | 0 | 0 |
| Bh226 | sexual | 0 | 1 |
| Bh227 | apomictic | 1 | 1 |
| Bh228 | apomictic | 1 | 0 |
| Bh229 | apomictic | 1 | 1 |
| Bh231 | sexual | 0 | 1 |
| Bh232 | apomictic | 1 | 1 |
| Bh233 | sexual | 0 | 1 |
| Bh234 | sexual | 0 | 0 |
| Bh235 | apomictic | 1 | 1 |
| Bh237 | sexual | 0 | 0 |
| Bh239 | sexual | 0 | 1 |
| Bh240 | apomictic | 1 | 1 |
| Bh241 | apomictic | 1 | 1 |
| Bh242 | apomictic | 1 | 0 |
| Bh243 | sexual | 0 | 0 |
| Bh244 | sexual | 0 | 1 |
| Bh245 | sexual | 0 | 0 |
| Bh246 | sexual | 0 | 0 |
| Bh247 | sexual | 0 | 0 |
| Bh248 | sexual | 0 | 0 |
| Bh249 | sexual | 0 | 0 |
| Bh250 | sexual | 0 | 0 |
| Bh252 | sexual | 0 | 0 |
| Bh254 | apomictic | 1 | 1 |
| Bh255 | apomictic | 1 | 1 |
| Bh256 | sexual | 0 | 0 |
| Bh257 | sexual | 0 | 1 |
| Bh258 | sexual | 0 | 1 |
| Bh259 | apomictic | 1 | 1 |
| Bh260 | apomictic | 1 | 1 |
| Bh261 | apomictic | 1 | 1 |
| Bh262 | sexual | 0 | 0 |
| Bh263 | sexual | 0 | 0 |
| Bh264 | apomictic | 1 | 0 |
| Bh265 | sexual | 0 | 0 |
| Bh267 | apomictic | 1 | 1 |
| Bh268 | apomictic | 1 | 1 |
| Bh269 | apomictic | 1 | 1 |
| Bh270 | apomictic | 1 | 1 |
| Bh272 | auto | - | 1 |
| Bh273 | apomictic | 1 | 1 |
| Bh274 | apomictic | 1 | 1 |
| Bh276 | apomictic | 1 | 1 |
| Bh277 | sexual | 0 | 0 |
| Bh278 | apomictic | 1 | 1 |
| Bh279 | apomictic | 1 | 1 |
| Bh281 | sexual | 0 | 0 |
| Bh283 | - | - | 0 |
| Bh284 | sexual | 0 | 0 |
| Bh285 | sexual | 0 | 1 |
| Bh287 | apomictic | 1 | 1 |
| Bh289 | sexual | 0 | 0 |
| Bh290 | sexual | 0 | 0 |
| Bh291 | sexual | 0 | 0 |
| Bh292 | sexual | 0 | 0 |
| Bh293 | sexual | 0 | 1 |
| Bh296 | sexual | 0 | 1 |
| Bh297 | apomictic | 1 | 1 |
| Bh302 | sexual | 0 | 0 |
| Bh303 | sexual | 0 | 0 |
| Bh309 | apomictic | 1 | 1 |
| Bh310 | apomictic | 1 | 1 |
| Bh311 | apomictic | 1 | 1 |
| Bh312 | - | - | 1 |
| Bh314 | sexual | 0 | 0 |
| Bh318 | sexual | 0 | 1 |
| Bh319 | apomictic | 1 | 1 |
| Bh320 | apomictic | 1 | 1 |
| Bh321 | sexual | 0 | 1 |
| Bh322 | apomictic | 1 | 0 |
| Bh323 | apomictic | 1 | 1 |
| Bh324 | apomictic | 1 | 1 |
| Bh325 | apomictic | 1 | 0 |
| Bh328 | sexual | 0 | 1 |
| Bh329 | sexual | 0 | 0 |
| Bh330 | apomictic | 1 | 1 |
| Bh331 | sexual | 0 | 0 |
| Bh333 | sexual | 0 | 0 |
| Bh334 | sexual | 0 | 0 |
| Bh335 | sexual | 0 | 0 |
| Bh336 | sexual | 0 | 1 |
| Bh337 | sexual | 0 | 0 |
| Bh339 | sexual | 0 | 0 |
| Bh340 | apomictic | 1 | 1 |
| Bh341 | sexual | 0 | 1 |
| Bh342 | sexual | 0 | 0 |
| Bh343 | sexual | 0 | 0 |
| Bh344 | sexual | 0 | 0 |
| Bh345 | sexual | 0 | 0 |
| Bh346 | sexual | 0 | 1 |
| Bh347 | apomictic | 1 | 1 |
| Bh348 | apomictic | 1 | 1 |
| Bh350 | sexual | 0 | 1 |
| Bh351 | apomictic | 1 | 1 |
| Bh352 | sexual | 0 | 1 |
| Bh353 | apomictic | 1 | 1 |
| Bh354 | sexual | 0 | 1 |
| Bh355 | sexual | 0 | 0 |
| Bh360 | sexual | 0 | 0 |
| Bh361 | sexual | 0 | 0 |
| Bh363 | sexual | 0 | 1 |
| Bh364 | sexual | 0 | 0 |
| H16 (female parent) | apomictic | 1 | 1 |
| H31 (male parent) | sexual | 0 | 0 |
| * The reproductive mode of each F_1_ individual was previously assessed by examining embryo sacs | | | |
| ** Presence (1) and absence (0) of bands | | | |

**Supplementary Alignment 1****:** The parents and hybrids of the mapping population were analyzed with p778/p779, a primer pair specific to the candidate gene for the parthenogenesis component of apomixis that has been developed for *Pennisetum squamulatum* (primer sequence: TATGTCACGACAAGAATATG; TGTAACCATAACTCTCAGC). Sanger technology was used to sequence the p778/779 amplicon of three randomly chosen apomictic hybrids (named H261, H310 and H330), in addition to four replicates of the apomictic parental genotype H016. Consensus singleton sequences were generated and compared to nr/nt, and the results showed significant blastn matches (E = 10-10) with ASGR-BBM-like1 gene sequences from *Cenchrus ciliaris* and *Pennisetum squamulatum*. Then, the singleton sequences were aligned with these gene sequences, and the multiple alignment revealed insertions and deletions (indels), in addition to SNPs (both colored in the alignment).

```
CLUSTAL O(1.2.4) multiple sequence alignment


consensus_apomictic_reverse             TGGAAGTTGAACTTATATATTTCACAAATGGATTGACATAGAACATATATTTGTGATACA	60
consensus_apomictic_foward_revcomp      ------------------------------------------------------------	0
EU559280.1                              TGGAAACTGAACTTATATATTTCACAAATGGATTGACATAGAACATATATTTGTGATACA	60
EU559278.1                              TGGAAACTGAACTTATATATTTCACAAATGGATTGACATAGAACATATATTTGTGATACA	60


consensus_apomictic_reverse             GGAAAAGCAGTGGTTTTTCTCGTGGTGCATCAATTTACCGAGGGGTTACCAGGTACAAAA	120
consensus_apomictic_foward_revcomp      ------------------------------------------------------------	0
EU559280.1                              GGAAAAGCAGTGGTTTTTCTCGTGGTGCATCAATTTACCGAGGGGTTACCAGGTACAAAA	120
EU559278.1                              GGAAAAGCAGTGGTTTTTCTCGTGGTGCATCAATTTACCGAGGGGTTACCAGGTACAAAA	120


consensus_apomictic_reverse             TATTCCTTTTCCTTATTATCATCTGGTTTTAGTTAGCAAGTGCATTGTTTCTATGGGAAT	180
consensus_apomictic_foward_revcomp      ------------------------------------------------------------	0
EU559280.1                              TATTCCTTTTCCTTATTATCTC-TGGTTTTAGTTAGCAAGTGCATTGTTTCTATGGGAAT	179
EU559278.1                              TATTCCTTTTCCTTATTATCTC-TGGTTTTAGTTAGCAAGTGCATTGTTTCTATGGGAAT	179


consensus_apomictic_reverse             TTGTG-----------------------TTGCACGTAGATCATAAATAGTTGCAACTATT	217
consensus_apomictic_foward_revcomp      ------------------------------------------------------------	0
EU559280.1                              TTGTGTTGCATGTAGATGGGAATTTGTGTTGCATGTAGATCATAAATAGTTGCAACTATT	239
EU559278.1                              TTGTGTTGCATGTAGATGGGAATTTGTGTTGCATGTAGATCATAAATAGTTGCAACTATT	239


consensus_apomictic_reverse             AATCTCATCG------TTCTGAATAGTTGTGGTACTCCTTTACCACAGTTGACTATGATA	271
consensus_apomictic_foward_revcomp      ------------------------------------------------------------	0
EU559280.1                              AATCTCATCGTTCTATTGCTGAATAGTTGTGGTACTCCTTTACCACAGTTGACTATGATA	299
EU559278.1                              AATCTCATCGTTCTATTGCTGAATAGTTGTGGTACTCCTTTACCACAGTTGACTATGATA	299


consensus_apomictic_reverse             TTCAATTATATTATTTTTCTTGCAAAGTTGATATTTAATTGCTTGTCTAGCTAACTTTCA	331
consensus_apomictic_foward_revcomp      ------------------------------------------------------------	0
EU559280.1                              TTCTATTATATTATTTTTCTTGCAAAGTTGATATTTAATTGCTTGTCTAGCTAACTTTCA	359
EU559278.1                              TTCTATTATATTATTTTTCTTGCAAAGTTGATATTTAATTGCTTGTCTAGCTAACTTTCA	359


consensus_apomictic_reverse             AGCA--------------------------------------------------------	335
consensus_apomictic_foward_revcomp      ------------------------------------------------------------	0
EU559280.1                              AGCAATCATGTAAAACAGGCACCATCAGCATGGAAGGTGGCAAGCAAGAATAGGAAGTGT	419
EU559278.1                              AGCAACCATGTAAAACAGGCACCATCAGCATGGAAGGTGGCAAGCAAGAATAGGAAGTGT	419


consensus_apomictic_reverse             ------------------------------------------------------------	335
consensus_apomictic_foward_revcomp      ------------------------------------------------------------	0
EU559280.1                              GGCAGGAAACAAGGATCTTTATTTGGGCACATTCAGTAAGTCACATTTTAATATTTTTAA	479
EU559278.1                              GGCAGGAAACAAGGATCTTTATTTGGGCACATTCAGTAAGTCACATTTTAATATTTTTAA	479


consensus_apomictic_reverse             ------------------------------------------------------------	335
consensus_apomictic_foward_revcomp      -----------------------CAAGCAAAATGGAAGCAAGACAGAAAAGCATAAACCT	37
EU559280.1                              TGAAGCACTGATTTTTTTT-TGTCAAGCAAAATGGAAGCAAGACAGAAAAACATAAACC-	537
EU559278.1                              TGAAGCACTGATTTTTTTTTTGTCAAGCAAAATGGAAGCAAGACAGAAAAACATAAACCT	539


consensus_apomictic_reverse             ------------------------------------------------------------	335
consensus_apomictic_foward_revcomp      ACTGCTGGAGCACCTTTTTCATTATTTTATCTCTTGAATATAATAGTATGTGGCTGACCT	97
EU559280.1                              --TACTGGAGCACCTTTTTCATTATTTTGTCTCTTGAATATAATAGTATGTGGCTGACCT	595
EU559278.1                              ACTGCTGGAGCACCTTTTTCATTATTTTGTCTCTTGAATATAATAGTATGTGGCTGACCT	599


consensus_apomictic_reverse             ------------------------------------------------------------	335
consensus_apomictic_foward_revcomp      CTCCCTGTGTAGGTACCCAGGAGGAAGCTGCAGAGGCTTACGACATTGCTGCCATCAAAT	157
EU559280.1                              CTCCCTGTGTAGGTACCCAGGAGGAAGCTGCAGAGGCTTACGACATTGCTGCCATCAAAT	655
EU559278.1                              CTCCCTGTGTAGGTACCCAGGAGGAAGCTGCAGAGGCTTACGACATTGCTGCCATCAAAT	659


consensus_apomictic_reverse             ------------------------------------------------------------	335
consensus_apomictic_foward_revcomp      TCCAAGGCCTCAATGCTGTCACGAACTTTGACATGAGCCGGTATGACGTCAAGAGCATCA	217
EU559280.1                              TCCGAGGCCTCAATGCTGTCACGAACTTTGACATGAGCCGGTATGACGTCAAGAGCATCA	715
EU559278.1                              TCCGAGGCCTCAATGCTGTCACGAACTTTGACATGAGCCGGTATGACGTCAAGAGCATCA	719


consensus_apomictic_reverse             ------------------------------------------------------------	335
consensus_apomictic_foward_revcomp      TTGAGAGCAGCTCCTTGCCTGTTGGCGGCACTCCAAAGCGTCTCAAGGAAGTGCCTGATC	277
EU559280.1                              TTGAGAGCAGCTCCCTGCCTGTTGGCGGCACTCCAAAGCGTCTCAAGGAAGTGCCTGATC	775
EU559278.1                              TTGAGAGCAGCTCCCTGCCTGTTGGCGGCGCTCCAAAGCGTCTCAAGGAAGTGCCTGATC	779


consensus_apomictic_reverse             ----------------------------------------------------	335
consensus_apomictic_foward_revcomp      AATCAGTGGGCATCAACATAAACGGTGCTGAGTCTGCTGGTCATATGACTGC	329
EU559280.1                              AATCAGATATGGGCATCAACATAAACGGTGACTCTGCTGGTCATATGACTGC	827
EU559278.1                              AATCAGATATGGGCATCAACATAAACGGTGACTCTGCTGGTCATATGACTGC	831
```
